# The P_5_-ATPase ATP13A1 modulates MR1-mediated antigen presentation

**DOI:** 10.1101/2021.05.26.445708

**Authors:** Corinna A. Kulicke, Erica De Zan, Zeynep Hein, Claudia Gonzalez-Lopez, Swapnil Ghanwat, Natacha Veerapen, Gurdyal S. Besra, Paul Klenerman, John C. Christianson, Sebastian Springer, Sebastian Nijman, Vincenzo Cerundolo, Mariolina Salio

**Affiliations:** MRC Human Immunology Unit, Weatherall Institute of Molecular Medicine, Radcliffe Department of Medicine, University of Oxford, Oxford, UK; Ludwig Institute for Cancer Research Ltd. and Target Discovery Institute, Nuffield Department of Medicine, University of Oxford, Oxford, UK; Department of Life Sciences and Chemistry, Jacobs University, Bremen, Germany; School of Biosciences, University of Birmingham, Birmingham B11 2TT, United Kingdom; Peter Medawar Building, Nuffield Department of Medicine, University of Oxford; Translational Gastroenterology Unit, Nuffield Department of Medicine, University of Oxford; Botnar Research Centre, Nuffield Department of Orthopaedics, Rheumatology and Musculoskeletal Sciences, University of Oxford, Oxford, UK

**Keywords:** MHC I-related protein 1 (MR1), mucosal-associated invariant T cell (MAIT), MR1-restricted T cell (MR1T), antigen presentation, protein trafficking, HAP1, gene trap, ATP13A1, P5-type ATPase

## Abstract

The monomorphic antigen presenting molecule MHC-I-related protein 1 (MR1) presents small molecule metabolites to mucosal-associated invariant T (MAIT) cells. The MR1-MAIT cell axis has been implicated in a variety of infectious and non-communicable diseases and recent studies have begun to develop an understanding of the molecular mechanisms underlying this specialised antigen presentation pathway. Yet, the proteins regulating MR1 folding, loading, stability, and surface expression remain to be identified. Here, we performed a gene trap screen to discover novel modulators of MR1 surface expression through insertional mutagenesis of an MR1-overexpressing clone derived from the near-haploid human cell line HAP1 (HAP1.MR1). The most significant positive regulators identified included β_2_-microglobulin, a known regulator of MR1 surface expression, and ATP13A1, a P_5_-ATPase in the endoplasmic reticulum (ER) with putative transporter function not previously associated with MR1-mediated antigen presentation. CRISPR/Cas9-mediated knock-out of ATP13A1 in both HAP1.MR1 and THP-1 cell lines revealed a profound reduction in MR1 protein levels and a concomitant functional defect specific to MR1-mediated antigen presentation. Collectively, these data are consistent with the ER-resident ATP13A1 as a key post-transcriptional determinant of MR1 surface expression.

## Introduction

The monomorphic major histocompatibility complex (MHC) class I-related protein 1 (MR1) presents small molecule metabolites to mucosal-associated invariant T (MAIT) cells (1–3). While the full spectrum of MR1 ligands is still being elucidated (2, 4–10), the best characterized MAIT-activating MR1 ligands are derivatives of pyrimidine intermediates of riboflavin biosynthesis, a pathway specific to certain fungi and bacteria and, thus, intrinsically non-self for humans (1, 11). Accordingly, MAIT cells are activated upon recognition of MR1 ligands derived from a variety of microbes capable of riboflavin synthesis such as *Escherichia coli* (*E.coli*), *Salmonella typhimurium*, and *Mycobacterium tuberculosis* (12–15). Molecularly, MAIT cells are characterized by high expression of the C-type lectin CD161, the interleukin (IL)-18 receptor, CD26 (16–18) and the canonical Vα7.2-Jα33/12/20 T cell receptor (TCR) α chain in humans (19–21). Owing to their restricted TCR repertoire, their relative abundance, and their effector-memory phenotype, MAIT cells are counted as an innate-like lymphocyte subset (22). Their high numbers and poised phenotype allow them to rapidly carry out effector functions in response to their cognate microbial antigens and place them at the interface between the innate and the adaptive immune system (22, 23). Furthermore, the MR1 antigen presentation pathway is an attractive target for both therapeutic and vaccination strategies as the monomorphic nature of the presenting molecule renders it independent of individual variations in human leukocyte antigen (HLA) expression (24–28). A better understanding of this non-classical antigen presentation pathway is, thus, crucial for harnessing the protective potential of MAIT cells in such therapeutic approaches.

MR1 transcript is detectable in almost all human cell lines and tissues but endogenous surface expression of MR1 protein is difficult to detect in most human cells even upon exposure to microbial MR1 ligand (29, 30). Nevertheless, such low MR1 surface levels are capable of inducing a potent interferon-γ (IFNγ) response in MAIT cells (29, 30). Since endogenous MR1 is difficult to detect, most of the current knowledge has been obtained in over-expression systems (30–35). In one widely used model of MR1 overexpression, the majority of cellular MR1 resides in the ER in a partially folded, ligand-receptive state in the absence of exogenous or microbial ligands (34, 36). Upon ligand binding, the MR1 heavy chain-β_2_-microglobulin (β_2_m)-complex translocates to the cell surface (34, 37). ER egress is thought to require neutralisation of the positive charge on the lysine 43 (K43) residue within the MR1 ligand binding groove (34, 37). In other studies, MR1 has been found to partially co-localise with LAMP1^+^ in endolysosomal compartments (35, 38) and the relative contributions of these two intracellular pools to antigen presentation under physiological conditions remain to be determined (39). Importantly, a growing body of evidence suggests that the loading and trafficking of MR1 in the context of intracellular bacterial infection differs from the presentation of exogenous ligand (30, 35, 39–42).

In light of the ubiquitous expression of MR1 transcript and the observation that activating MR1 ligands can be derived from both pathogenic and commensal microorganisms, tight regulation of this highly sensitive antigen presentation pathway is required (31, 41). Using a previously described functional genetic screen utilising the near-haploid human cell line HAP1 (43, 44), we identified the P_5_-ATPase ATP13A1 as one of the proteins involved in the modulation of MR1 surface expression.

## Results

### A gene trap screen identifies the P_5_-ATPase ATP13A1 as a putative modulator of MR1 surface expression

The near-haploid human cell line HAP1 is a powerful tool for genetic loss-of-function screens as only one copy of a gene needs to be mutated to achieve a functional knock-out (43). Since HAP1 WT cells did not express appreciable amounts of MR1 at the protein or transcript level (Supporting Information Figure 1), we transduced HAP1 cells with lentiviral particles encoding the MR1 complementary DNA (cDNA) sequence. To ensure that any difference in MR1 surface levels observed in the screen was due to gene editing rather than varying copy numbers of the MR1 cassette in the polyclonal HAP1.MR1 population, we single cell sorted MR1^+^ cells (Supporting Information Figure 1). The resulting HAP1.MR1 parent clones were analysed for ploidy and responsiveness to the MR1-stabilising ligand Acetyl-6-formylpterin (45) (Ac6FP) (Supporting Information Figure 1). The HAP1.MR1 parent clone D9 was expanded and transduced with a gene trap virus as previously described (44) (see Experimental Procedures for details). Subsequently, the polyclonal mutant population was pulsed with Ac6FP to induce MR1 surface translocation and surface stained for MR1. We then sorted the MR1^hi^ and MR1^low^ tails of the distribution by flow cytometry (Supporting Information Figure 2) and sequenced the DNA to identify the retroviral insertion sites (44). In total, >1.75×10^6^ unique insertion sites were recovered for each sorted population with approximately 48% being sense insertions (Supporting Information Table 1). The sense insertions across both sorted populations mapped to 16,952 genes, covering about 85% of the currently predicted protein-coding genes in the human genome (46). Out of these, 199 genes were scored as putative regulators of MR1 surface expression (false discovery rate (FDR)-corrected p-value (fcpv) < 0.01), including the MR1 transgene itself and β_2_m (Figure 1, Supporting Information Table 2). Intriguingly, putative negative regulators included HLA-A as well as components of the peptide loading complex (PLC; TAP1, TAP2, and TAPBP, the gene encoding tapasin (47)) but also genes involved in the regulation of protein transport (e.g. TMEM131 (48) and RALGAPB (49)), endolysosomal trafficking and homeostasis (e.g. LAMTOR2 (50, 51) and VAC14 (52, 53)), N-terminal protein acetylation (e.g. NAA30 (54)), and ubiquitination (e.g. CUL3 (55, 56)) (all highlighted in Figure 1). Similarly, the putative positive regulators comprised genes with diverse functions ranging from implication in immune responses (e.g. B2M (57, 58) and IRF2 (59, 60)) to endosomal recycling (e.g. VPS29 (61) and VPS53 (62)), cytoskeletal organisation (e.g. NHLRC2 (63, 64)), glycosylation (e.g. GANAB (65) and SPPL3 (66)), mRNA processing (e.g. PARN (67) and ZCCHC14 (68)), and ion homeostasis in the ER (ATP13A1 (69)) (all highlighted in Figure 1).

**Figure 1:**
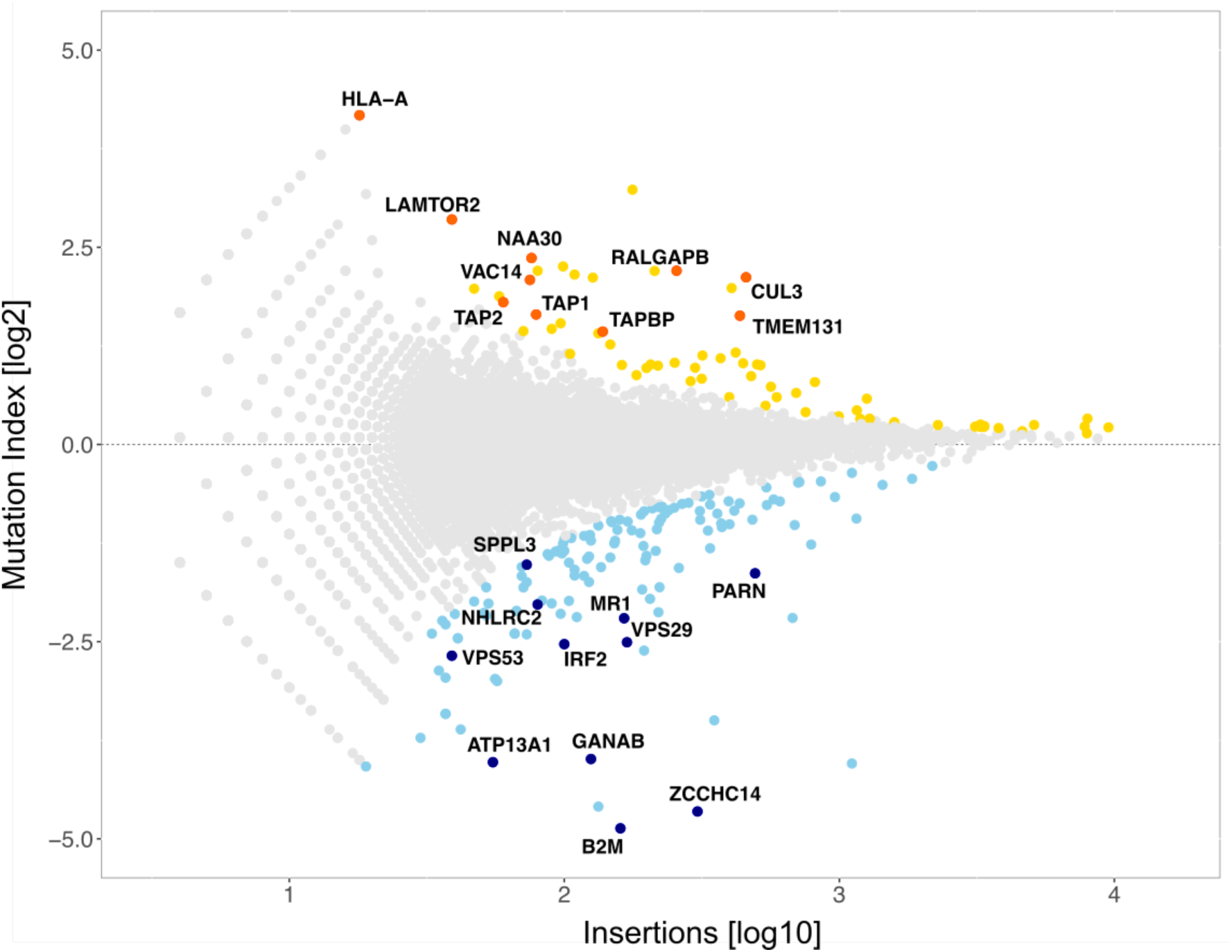
A gene-trap screen in HAP1.MR1 clone D9 identifies putative positive and negative modulators of MR1 surface expression. For each gene, the odds ratio of it being enriched in the MR1^high^ compared to the MR1^low^ population was calculated and plotted as the mutation index (MI) against the total number of sense insertions (see Methods). Genes significantly (fcpv < 0.01) enriched in the high or low fraction are highlighted in yellow and blue, respectively, and represent putative negative and positive regulators of MR1 surface expression. MR1, its known regulator β2m (encoded by gene B2M), and selected genes based on functional annotations are highlighted (see text). Raw data are available in Supporting Information Table 2.

The difference in the number of positive and negative candidate genes identified reflects the fact that the screen setup was more powerful in detecting putative positive regulators than negative ones. The screen was carried out in the presence of Ac6FP which results in the translocation of most MR1 molecules to the surface. Thus, a further increase in MR1 surface expression, as would be expected upon knock-out of a negative regulator, may be masked. Loss of a positive regulator, on the other hand, leads to reduced MR1 surface levels and was more easily detected in our system. Therefore, we focused our analysis on the putative positive regulators of MR1 surface expression.

The most highly enriched gene in the MR1^low^ population (log2MI = −4.867, fcpv = 1.47×10^-34^) was B2M, the gene encoding β_2_ – microglobulin (β_2_m_;_ Figure 1, Supporting Information Table 2). Expression of β_2_m is a known requirement for MR1 surface translocation (32, 70) and can, thus, be considered a positive control for the screen. Based on functional annotations available in public databases including Pubmed Gene, GeneCards (71), and UniProtKB (72), we chose a subset of the identified putative regulators for further validation and characterization. Among those, ZCCHC14 (log2MI = −4.653, fcpv = 3.10×10^-65^), ATP13A1 (log2MI = −4.028, fcpv = 2.28×10^-9^), GANAB (log2MI = −3.988, fcpv = 7.46×10^-23^), VPS53 (log2MI = −2.678, fcpv = 4.55×10^-4^), and IRF2 (log2MI = −2.532, fcpv = 2.39×10^-10^) displayed an MR1 phenotype with at least one of two single guide RNAs (sgRNAs) targeting the gene in an initial validation in bulk HAP1.MR1 cells (Supporting Information Figure 3). Interestingly, targeting of VPS53 and IRF2 reduced MR1 surface levels at baseline but not in the presence of Ac6FP (Supporting Information Figure 3), indicating that these modulators are less important for MR1 surface expression when MR1 ligand is available to fold and stabilize the molecule.

One of the most prominent and functionally interesting putative positive regulators of MR1 surface expression was the P_5_-ATPase ATP13A1 (Figure 1, Supporting Information Table 2). While information on mammalian ATP13A1 has been sparse, recent evidence points towards a dislocase function (73). The yeast homologue Spf1p (also known as Cod1p) localises to the ER and is involved in ion homeostasis (69, 74). Consequently, loss of *spf1* causes hypoglycosylation of proteins and results in ER stress (69, 74–78). In the context of MR1 antigen presentation, ATP13A1 was a particular promising target because of its localisation to the ER and its putative transporter function.

### ATP13A1 modulates the size of the cellular MR1 pool

To investigate the role of ATP13A1 in MR1-mediated antigen presentation we generated HAP1.MR1 ATP13A1 knock-out (KO) clones. The same HAP1.MR1 parent clone used in the original screen (clone D9) was transiently transfected with plasmids encoding the Cas9 protein, the fluorescent protein mRuby, and one of three different sgRNAs targeting different regions of the ATP13A1 gene (Supporting Information Figure 4). Single mRuby^+^ cells were sorted and individual clones screened for loss of ATP13A1 expression by Western blot (Figure 2*A*). Strikingly, total cellular MR1 levels were reduced in all but one HAP1.MR1 ATP13A1 KO clones regardless of the sgRNA used (Figure 2*A* and Supporting Information Figure 5). A notable exception was clone 3-15 which displayed reduced cellular MR1 levels despite expressing ATP13A1 (see below and Supporting Information Figure 5). Sequencing revealed that a six base pair deletion in clone 3-15 left the reading frame of ATP13A1 intact but altered a highly conserved region within the amino acid sequence (79) (Supporting Information Figure 6). The importance of this GxPF sequence is not only underscored by its conservation across yeast, murine and human ATP13A1 homologues (75, 79, 80) (NCBI Reference Sequences NP_010883.3, NP_573487.2, and NP_065143.2, respectively), but also by the observation that a construct lacking part of this motif was unable to rescue the *spf1* KO phenotype in yeast (75). Alterations to this region appear to have resulted in the production of a non-or dys-functional protein, which could explain the “KO-like” phenotype of clone 3-15 (Supporting Information Figure 5). This makes clone 3-15 a very interesting control as it suggests that loss of ATP13A1 function and not merely expression of the protein is responsible for the MR1 phenotype.

**Figure 2:**
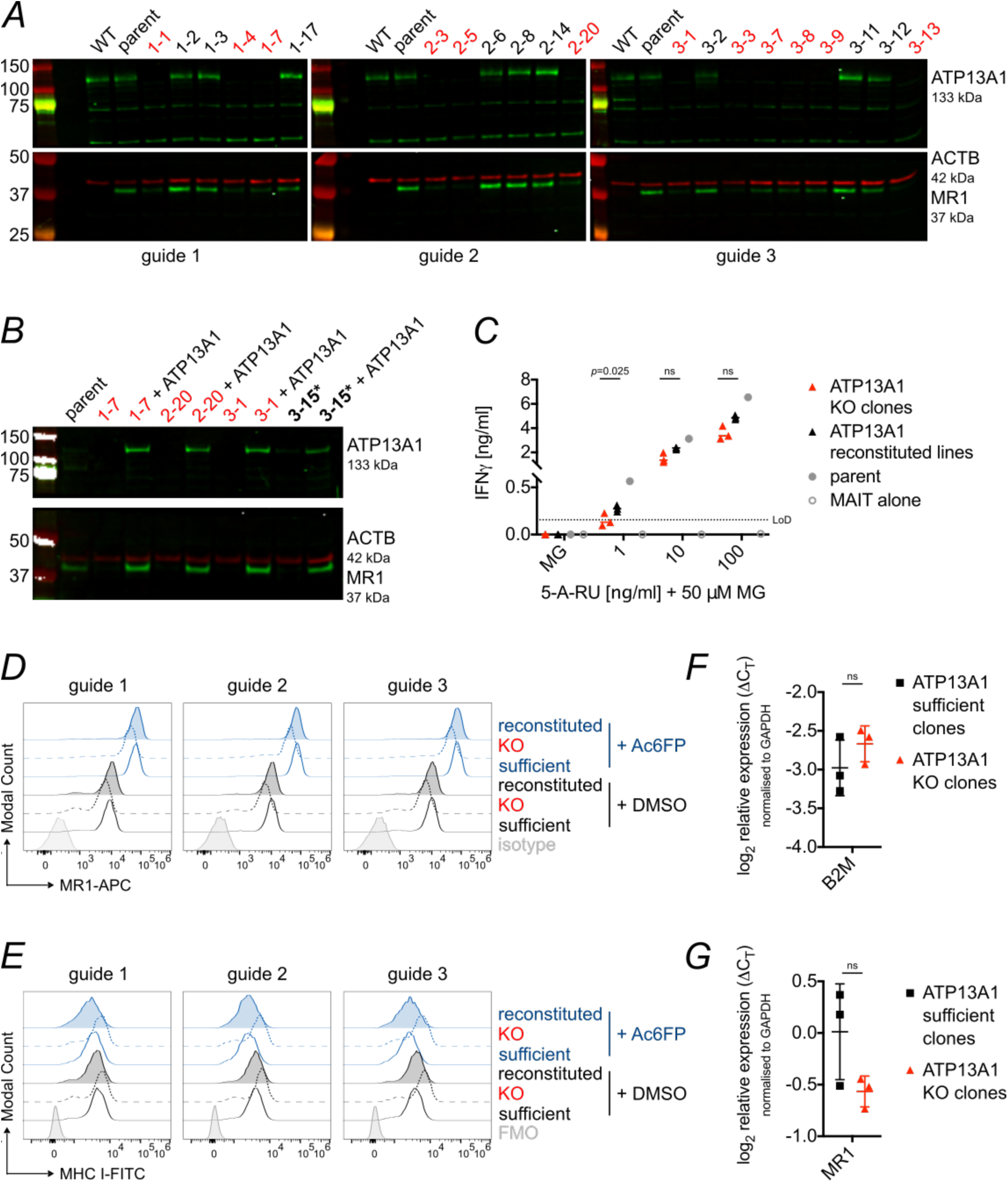
Cellular MR1 levels are reduced in HAP1.MR1 ATP13A1 KO clones. *A*, the HAP1.MR1 parent clone was transfected with CRISPR/Cas9 plasmids encoding three different sgRNAs targeting ATP13A1. Single cells were sorted by flow cytometry and generated clones were analysed by Western blot. Membranes were cut and probed with anti-ATP13A1 (top, green) or anti-MR1 (bottom, green) and anti-β actin (ACTB; bottom, red). Molecular weight of marker bands is shown in kDa to the left of the blots. Results are representative of at least two experiments for most clones. This includes the blot shown in Supporting Information Figure 5. *B*, ATP13A1 expression was reconstituted in a subset of clones using a lentiviral expression system and MR1 expression in the polyclonal transduced cell lines was analysed by Western Blot. *C*, the HAP1.MR1 parent clone, these KO clones and the reconstituted cell lines were pulsed with 5-A-RU+50 µM MG and incubated with sorted human MAIT cells for at least 36 h. IFNγ in the supernatants was quantified by ELISA. Each symbol represents the mean of technical duplicates for one clone per cell line and median is shown for the groups. Statistical significance of differences between ATP13A1 KO clones and reconstituted lines was analysed using two-tailed, paired *t* tests. The HAP1.MR1 parent clone D9 is shown for reference but was not included in statistical analyses. *D* + *E*, ATP13A1 sufficient (solid lines), KO (dashed lines), and reconstituted (filled) cells were incubated with 5 µg/ml of Acetyl-6-formylpterin (Ac6FP, blue) or DMSO (black) for 5 h before staining for MR1 (*D*) and MHC class I (*E*) surface expression. Isotype/fluorescence minus one (FMO) controls are from one sufficient, one KO, and one reconstituted clone each. *F* + *G*, B2M and MR1 transcript levels were compared in three ATP13A1 sufficient clones and three ATP13A1 KO clones. Each data point represents the mean of technical triplicates for one clone and mean and SD are shown for each group. Statistical significance of differences in transcript levels were analysed using an unpaired *t* test. Data in *C* + *D* are representative of three experiments for all clones except clone 3-1 and its reconstituted cell line although statistical significance varied across repeats (see Supporting Information Figure 7). Data in *F* + *G* are representative of two experiments. For HAP1 clones shown in *C*-*G* see Supporting Information Table 3. KO clones are highlighted in red. ns = *p* > 0.05. LoD = limit of detection.

When ATP13A1 expression was restored in a subset of clones comprising one KO clone for each sgRNA and clone 3-15, total MR1 levels were rescued, confirming that this phenotype is indeed attributable to loss of ATP13A1 and that the mutation found in clone 3-15 is not dominant negative (Figure 2*B*). Testing the same three HAP1.MR1 ATP13A1 KO clones in T cell stimulation assays, we observed a reduction in MAIT cell activation compared to the parental clone (Figure 2*C*). Importantly, ATP13A1 over-expression increased the antigen presentation capacity of all three HAP1.MR1 ATP13A1 KO clones and this trend was statistically significant at low antigen concentrations (Figure 2*C* and Supporting Information Figure 7). Although statistical significance varied across experimental replicates, the trends reported here were consistent across sgRNAs used for knock-out, MAIT cell donors, and assays, together strengthening the general conclusion that the MR1 phenotype can be rescued (at least partially) by reconstitution of ATP13A1 expression.

Next, we compared one clone per sgRNA that was deficient for ATP13A1 (“ATP13A1-deficient clones”) to one clone per sgRNA that was mRuby^+^ upon sorting but still expressed the protein (“ATP13A1-sufficient clones”). Details on editing of the clones used for the individual experiments can be found in Supporting Information Table 3 and Supporting Information Figure 6. Consistent with the results from the gene trap screen, surface levels of folded MR1 in HAP1.MR1 clones deficient for ATP13A1 were lower than those in their ATP13A1-sufficient counterparts both in the presence and absence of Ac6FP and independent of the sgRNA used to knock out ATP13A1 (Figure 2*D*). As with the total protein content, the reduced MR1 surface expression could be reversed by over-expression of ATP13A1 (Figure 2*D*). Importantly, surface expression of classical MHC class I molecules was not similarly impacted by loss of ATP13A1, suggesting that the phenotype is specific to MR1 (Figure 2*E*). Indeed, MHC class I surface levels appeared to be elevated in the ATP13A1 KO clones, possibly because of increased availability of β_2_m.

HAP1 cell cultures are prone to becoming polyploid over time, due to a competitive growth disadvantage of haploid cells (81). The flow cytometry and the functional data in Figure 2 are representative of two experiments, but one HAP1.MR1 ATP13A1 KO clone (3–1) lost its MR1 phenotype in a third repeat (Supporting Information Figure 7). This correlated with morphological changes indicative of a loss of the haploid state and so for further validation experiments we used THP-1 cells as a model system (see below).

Importantly, β_2_m was not differentially expressed in the KO compared to the sufficient clones at the transcript (Figure 2*F*) or protein level (Supporting Information Figure 8), indicating that the difference in MR1 surface expression was likely not due to a lack of its smaller subunit. This conclusion is corroborated by the rescue of both total and surface levels of MR1 in the reconstituted HAP1.MR1 ATP13A1 KO clones (Figure 2*B* and *D*) which demonstrates that β_2_m is not a limiting factor for MR1 expression in these cells. However, there was a trend towards lower MR1 transcript levels in the ATP13A1 KO clones (Figure 2*G*). Thus, we cannot exclude an effect of ATP13A1 on MR1 mRNA stability in the HAP1 system.

To confirm that the MR1 phenotype was not specific to the HAP1.MR1 parent clone used in the screen, we used sgRNA2 to knock out ATP13A1 in two other HAP1.MR1 parent clones (D3 and F2, compare Supporting Information Figure 1). These newly derived clones replicated the loss of total MR1 by Western blot and largely also the MR1 surface phenotype except for one of the three tested F2 ATP13A1 KO clones, which had likely become polyploid (Supporting Information Figures 9 and 10). Overall, these data indicate that the effect of ATP13A1 is not related to the number or site of insertions of the lentiviral MR1 expression cassette.

### ATP13A1 specifically affects MR1-mediated antigen presentation but not classical peptide presentation

Next, we validated the role of ATP13A1 in the monocytic cell line THP-1, which is frequently used as a model system for MR1-mediated antigen presentation (2, 30, 82, 83). We generated ATP13A1 KO clones from a THP-1 parent clone derived by limiting dilution (Figure 3*A*) and confirmed disruption of ATP13A1 on the genomic level by Next Generation Sequencing. This allowed us to investigate whether ATP13A1 deficiency impacted endogenous MR1. While MR1 mRNA levels were unaffected (Figure 3*B*), MR1 surface expression was reduced in five THP-1 ATP13A1 KO clones compared to ATP13A1-sufficient controls (Figure 3*C*). This trend did not reach statistical significance in one repeat due to one sample having high isotype staining (Supporting Information Figure 11). Importantly, endogenous MR1 surface levels were lower in ATP13A1 KO clones not only at baseline but also in the presence of the MR1-stabilising ligand Ac6FP (Figure 3*C*). Indeed, the fold-change of MR1 geometric mean fluorescence intensity (GeoMean) induced by Ac6FP was smaller in these clones (Figure 3*E*, top panel and Supporting Information Figure 11*C*, top panel), suggesting that the observed differences in MR1 surface expression are not simply due to the smaller pool of MR1 molecules available. While surface expression of classical MHC class I molecules was more heterogeneous than MR1 in ATP13A1 KO clones, there was no statistically significant difference compared to ATP13A1-sufficent clones (Figure 3*D*). As expected, MHC class I surface levels were unaffected by incubation with Ac6FP (Figure 3*E*, bottom panel).

**Figure 3:**
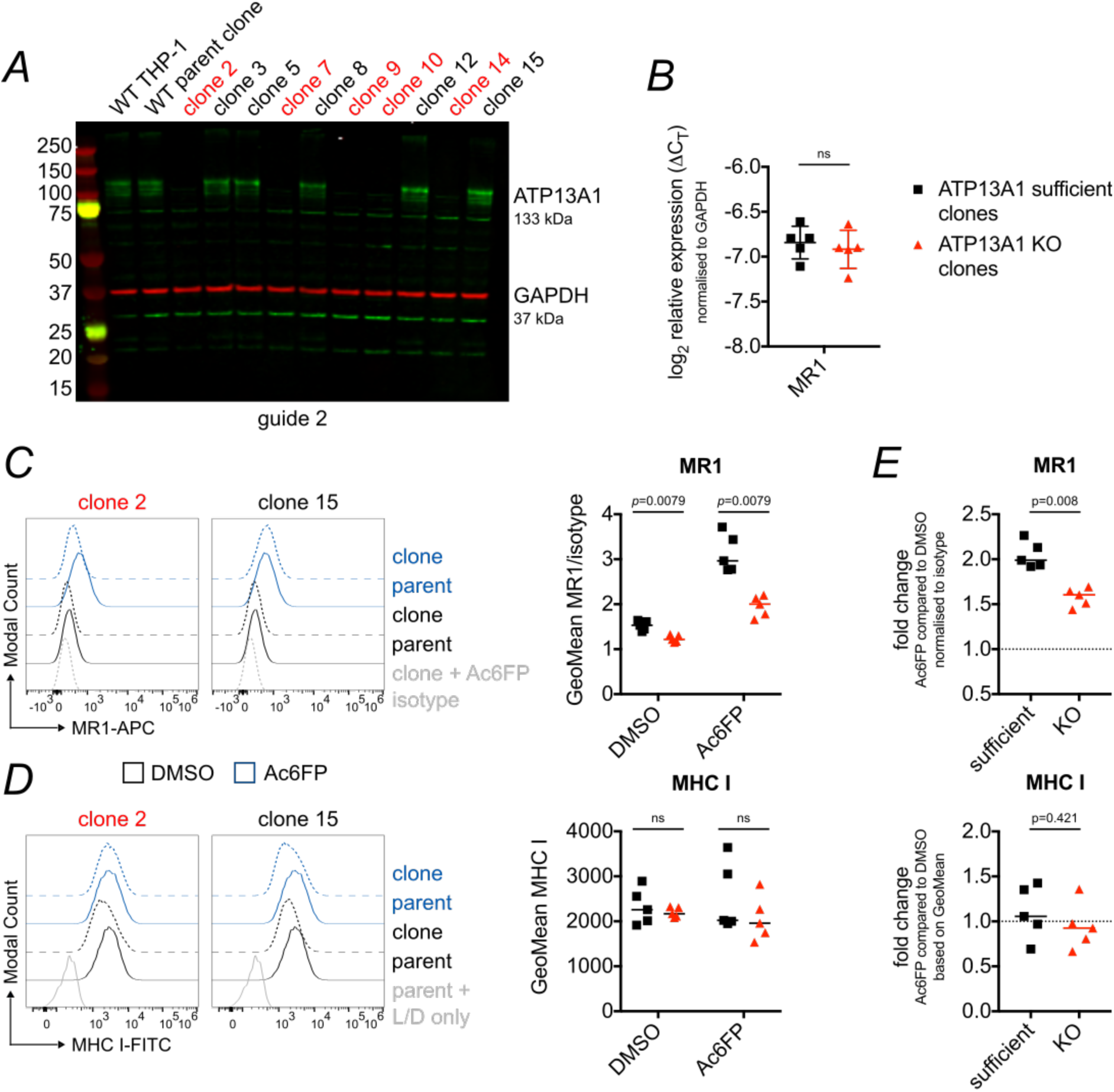
MR1 surface expression is reduced in THP-1 ATP13A1 KO clones. A THP-1 WT clone was transiently transfected with a CRISPR/Cas9 plasmid encoding sgRNA 2 targeting the first exon of ATP13A1 and cells were sorted into single wells. *A*, clones were screened by Western Blot. Membranes were probed with anti-ATP13A1 (green) and anti-β actin (ACTB; red) antibodies. Molecular weight of marker bands is shown in kDa to the left of the blot. *B*, MR1 transcript levels were compared in five ATP13A1 sufficient clones and five ATP13A1 KO clones. *C* + *D*, those ten clones were incubated with Acetyl-6-formylpterin (Ac6FP, blue) or DMSO (black) for 5 h before staining for MR1 (*C*) and MHC class I (*D*) surface expression. Histograms for two representative clones are shown on the left and cumulative data from all 10 clones are shown on the right. The same parent sample is shown in both histograms for comparison. *E*, the same data as in *C* + *D* shown as fold-change of Geometric Mean Fluorescence Intensity (GeoMean) with Ac6FP compared to DMSO control. Data in *A* are representative of at least three experiments for each clone except for WT THP-1 which was not always included. Results in *B*-*E* are supported by another experiment each although the repeat for *C* differed in statistical significance (see Supplementary Information Figure 11). Data are shown as mean with standard deviation in *B* and median in *C* - *E*. Statistical significance was calculated using a two-tailed *t*-test in *B* and the Mann-Whitney test in *C* - *E*. L/D = live/dead stain. ATP13A1 KO clones are highlighted in red. ns = *p* > 0.05.

When ATP13A1 expression was reconstituted in the THP-1 ATP13A1 KO clone 2, MR1 surface expression was restored (Supporting Information Figure 12). Total MR1 protein expression could not be investigated in these clones as endogenous MR1 levels are undetectable by Western blot (see e.g. Supporting Information Figure 12*B*).

Consistent with reduced levels of MR1 at the cell surface, the THP-1 ATP13A1 KO clones presented the MR1 ligand 5-(2-oxopropylideneamino)-6-D-ribitylaminouracil (5-OP-RU; added as 5-amino-ribityl uracil (5-A-RU) + methylglyoxal (MG)) less efficiently to sorted human MAIT cells than their ATP13A1 sufficient counterparts (Figure 4*A*). This trend was observed across multiple experiments using different MAIT cell donors and 5-A-RU concentrations, although it did not always reach statistical significance (Supporting Information Figure 13*A*). Since the antigen presentation pathways for exogenous ligand and intracellular bacteria differ mechanistically (31, 39, 42), we also tested the effect of ATP13A1 deficiency on MAIT cell activation in the context of bacterial infection. As with exogenous synthetic ligand, THP-1 ATP13A1 KO clones infected with *E.coli* showed a trend of eliciting a lower IFNγ response from sorted human MAIT cells as compared to ATP13A1 sufficient clones (Figure 4*B* and Supporting Information Figure 13*B*). Importantly, this functional defect appeared to be specific to MR1-mediated antigen presentation since the response of a peptide-specific T cell line to peptide-pulsed ATP13A1 KO clones was comparable to that elicited by ATP13A1 sufficient clones (Figure 4*C* and Supporting Information Figure 13*C*). Similarly, presentation of the CD1d ligand α-Galactosylceramide (αGalCer) to sorted human invariant natural killer T (iNKT) cells was not significantly affected by loss of ATP13A1 (Figure 4*D* and Supporting Information Figure 13*D*). Of note, ATP13A1 was not a hit in a recent whole-genome siRNA screen for modulators of antigen presentation by CD1d (84).

**Figure 4:**
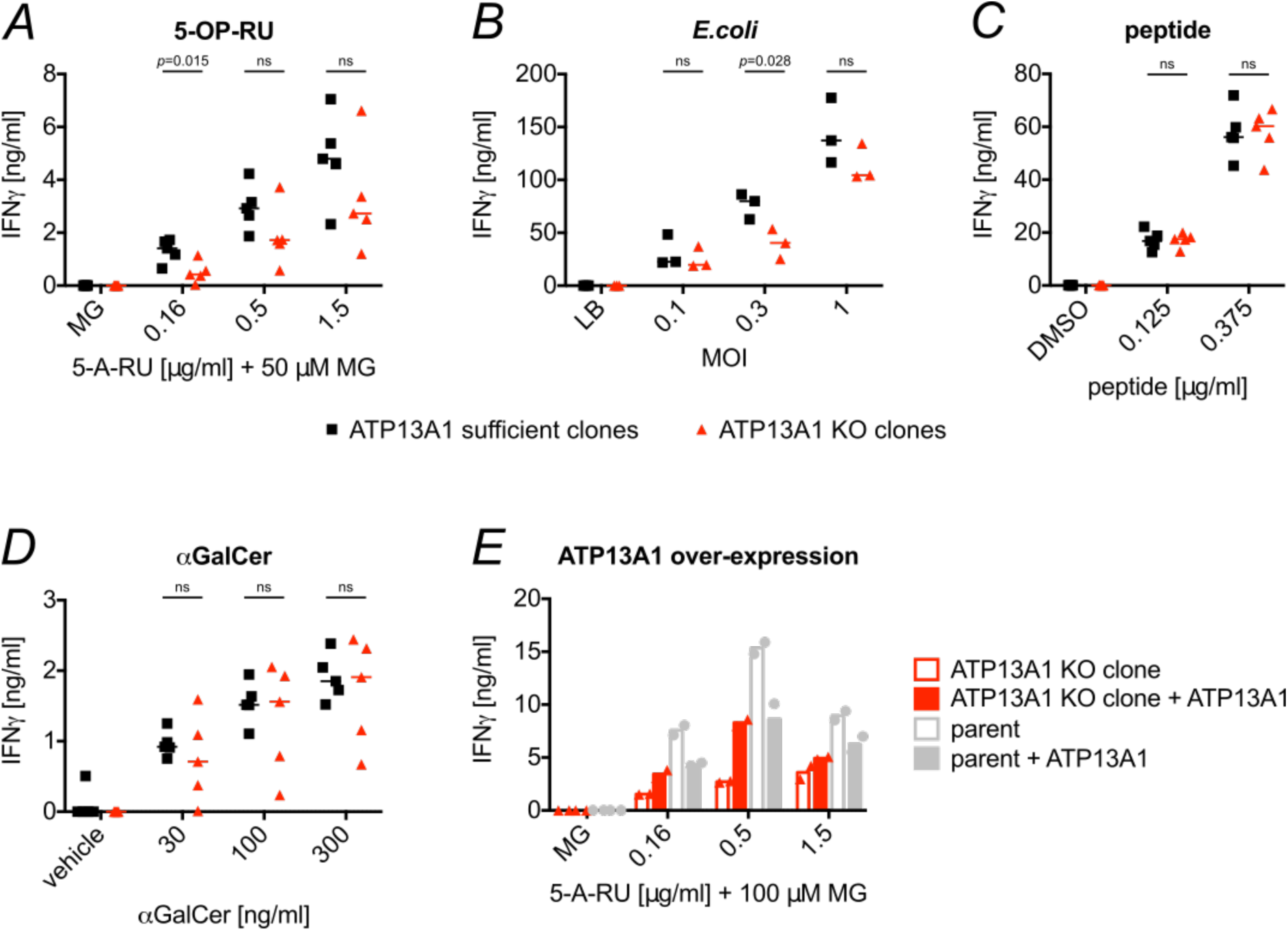
MR1-mediated antigen presentation is reduced in THP-1 ATP13A1 KO clones. *A* – *D*, THP-1-derived clones were pulsed with 5-A-RU+50 µM MG (*A*), *E. coli* (*B*), Melan A peptide (*C*), or αGalCer (*D*) and incubated with sorted human MAIT cells (5-A-RU, *E. coli*), sorted human iNKT cells (αGalCer), or a Melan A-specific T cell clone (peptide) for at least 36 h. *E*, The THP-1 parent clone and one THP-1 ATP13A1 KO clone were transduced with lentiviral particles encoding human ATP13A1. Transduced cells were pulsed with 5-A-RU+100 µM MG and incubated with sorted human MAIT cells for at least 36 h. IFNγ in the supernatants was quantified by ELISA. Each symbol represents the mean of technical duplicates for one clone and median is shown for the groups in *A* – *D*. Symbols in *E* represent mean of technical duplicates for two different MAIT cell donors tested in the same experiment. ATP13A1 KO clones are highlighted in red. Data are representative of at least two similar experiments each using different MAIT and iNKT cell donors where applicable although statistical significance varied between repeats (see Supporting Information Figure 13). Statistical significance of differences between indicated groups was analysed using unpaired *t* tests. ns = *p* > 0.05.

Importantly, when ATP13A1 expression was restored in ATP13A1 KO clone 2, its capacity to present 5-OP-RU was partially restored (Figure 4*E* and Supporting Information Figure 13*E*). Over-expression of ATP13A1 in the THP-1 parent clone, however, did not increase antigen presentation. Thus, it appears plausible that the transporter operates within a tightly controlled regulatory network optimised to prevent aberrant MAIT cell activation.

### ATP13A1-deficient HAP1.MR1 clones do not suffer from elevated ER stress

Having established that ATP13A1 KO clones display an antigen presentation phenotype largely specific to MR1 and attributable to ATP13A1 deficiency, we next investigated the mechanism by which ATP13A1 expression modulates MR1 protein levels. Since ATP13A1 has been implicated in the transport of cations (69, 74), we hypothesized that it could indirectly influence MR1 protein folding, loading, or stability in the ER by modulating ion concentrations in this compartment. Consistent with this, a large body of literature has linked deficiency for Spf1p and ATP13A1 to ER stress in yeast and human cells, respectively (69, 74, 76–78). We, thus, tested whether HAP1.MR1 clones deficient for ATP13A1 differentially up-regulated components of the unfolded protein response (UPR). Although all tested clones responded with up-regulation of the ER stress markers HERPUD1 (85) and ATF4 (86) when treated with the ER stress-inducing inhibitor thapsigargin (TG) (87), we found no evidence for differential UPR induction in the ATP13A1 KO clones (Figure 5*A* and *B*). The same pattern was evident when analysing the splice variants of XBP1, a transcription factor that controls the expression of UPR proteins in response to ER stress and is itself activated by alternative splicing of its mRNA (88) (Figure 5*C*). The data also indicate that the HAP1.MR1 ATP13A1 KO clones do not suffer from elevated ER stress at baseline, suggesting that the lower MR1 protein levels are not due to a general defect in protein folding.

**Figure 5:**
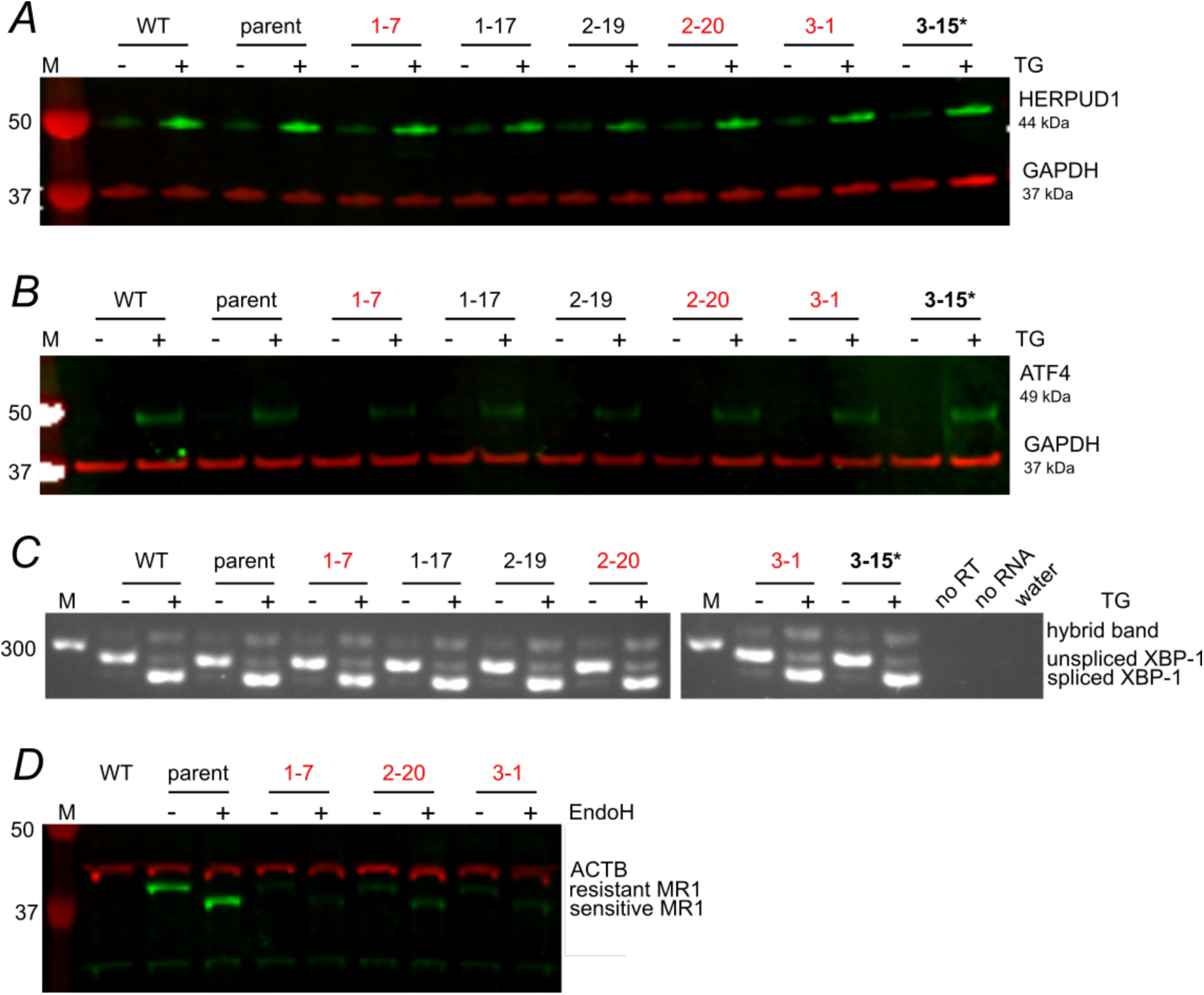
ATP13A1 knock out does not cause elevated ER stress in HAP1.MR1 ATP13A1 KO clones. *A* – *C*, HAP1.MR1-derived clones were treated with 0.1 µM thapsigargin (TG, +) or DMSO (-) for 3 h. Protein lysates were separated by SDS-PAGE and immuno-blotted for HERPUD1 (green) and loading control GAPDH (red) (*A*) or ATF4 (green) and GAPDH (red) (*B*). *C*, in the same experiment, cDNA of indicated HAP1.MR1 clones was analysed for the XBP1 splice variant on a 2.5% agarose gel. *D*, cell lysates of the indicated HAP1 clones were digested with endoglycosylase H (EndoH, +) or not (-). Digests were separated by SDS-PAGE and immuno-blotted for MR1 (green) and loading control ACTB (red). MR1-deficient HAP1 wild type (WT) lysates were included to identify unspecific binding. M denotes molecular weight or fragment size marker and values of visible marker bands are indicated in base pairs (bp) or kDa to the left of the images. Parent denotes HAP1.MR1 parent clone D9. Data in *A* - *C* are representative of two independent experiments except for the HAP1 WT which was not included in the repeat. Data in *D* are representative of four similar independent experiments including one that used the same lysates as the Western blot in Supporting Information Figure 8. ATP13A1 sufficient clones are different from Figure 2 (see Supporting Information Table 3). ATP13A1 KO clones are highlighted in red and the ATP13A1 mutant clone 3-15 is highlighted in bold and starred. noRT = no reverse transcriptase control.

Divalent cations such as calcium and manganese are important co-factors for many enzymes and disturbing their homeostasis leads to defects in glycoprotein processing (69, 74–76). Since the MR1 protein sequence features one known N-linked glycosylation site at Asn86 (32), we next determined whether MR1 in the HAP1.MR1 KO clones might be hypoglycosylated. To investigate the glycosylation state of MR1 in the HAP1.MR1 ATP13A1 KO clones, whole cell lysates were digested with Endoglycosidase H (EndoH). Since EndoH cannot cleave the highly complex polysaccharide residues generated during post-translational modification in the Golgi apparatus, sensitivity to EndoH digestion can also be used as a proxy of ER-residency (89, 90). It has previously been shown that in many cell lines MR1 primarily resides in a pre-Golgi compartment at steady state as reflected by its sensitivity to digestion with EndoH (32, 34, 70). Accordingly, the cellular MR1 pool in the HAP1.MR1 parent clone D9 was also sensitive to digestion with EndoH (Figure 5*D*). Interestingly, the much smaller pool of MR1 molecules in the HAP1.MR1 ATP13A1 KO clones was equally sensitive to the endoglycosidase, indicating that these molecules are glycosylated normally within the ER. This makes it seem unlikely that a general dysfunction of the ER glycosylation machinery is the cause for the reduced MR1 protein levels at baseline. Importantly, protein levels of classical MHC class I molecules were unaffected in the HAP1.MR1 ATP13A1 KO clones (Supporting Information Figure 14), further corroborating this conclusion. Of note, the antibody used to detect MHC class I by Western Blot does not react with HLA-A and HLA-G alleles (91). Hence, potential allele-specific effects on MHC class I molecules would not be detected in this assay and can, thus, not be excluded.

### ATP13A1 deficiency affects MR1 levels at or immediately after protein synthesis

Since MR1 transcript levels did not significantly differ between ATP13A1 KO and sufficient HAP1.MR1 (Figure 2*G*) or THP-1 (Figure 3*B*) clones and we found no evidence for a differential global induction of the unfolded protein response in the HAP1.MR1 ATP13A1 KO clones (Figure 5), we hypothesized that MR1 may be specifically targeted for ER-associated degradation (ERAD) in these cells. To test this hypothesis, we inhibited proteasomal degradation with the reversible peptide aldehyde inhibitor MG-132 (92, 93). In parallel, we inhibited ERAD at a different step with the irreversible p97 inhibitor NMS-873 (94). p97 aids in the extraction of ubiquitinated target proteins from the ER lumen into the cytosol and, thus, encourages misfolded proteins towards proteolysis (94, 95). Even though proteasomal degradation was successfully inhibited in this experiment as shown by the stabilisation of HERPUD1, this inhibition did not lead to an increase in full-length MR1 protein levels (Figure 6*A*). This suggests that MR1 is not differentially degraded by ERAD upon ATP13A1 KO under homeostatic conditions. However, proteasome inhibition may have led to the accumulation of ubiquitinated, high molecular weight MR1 species which could not be detected with the experimental setup used.

**Figure 6:**
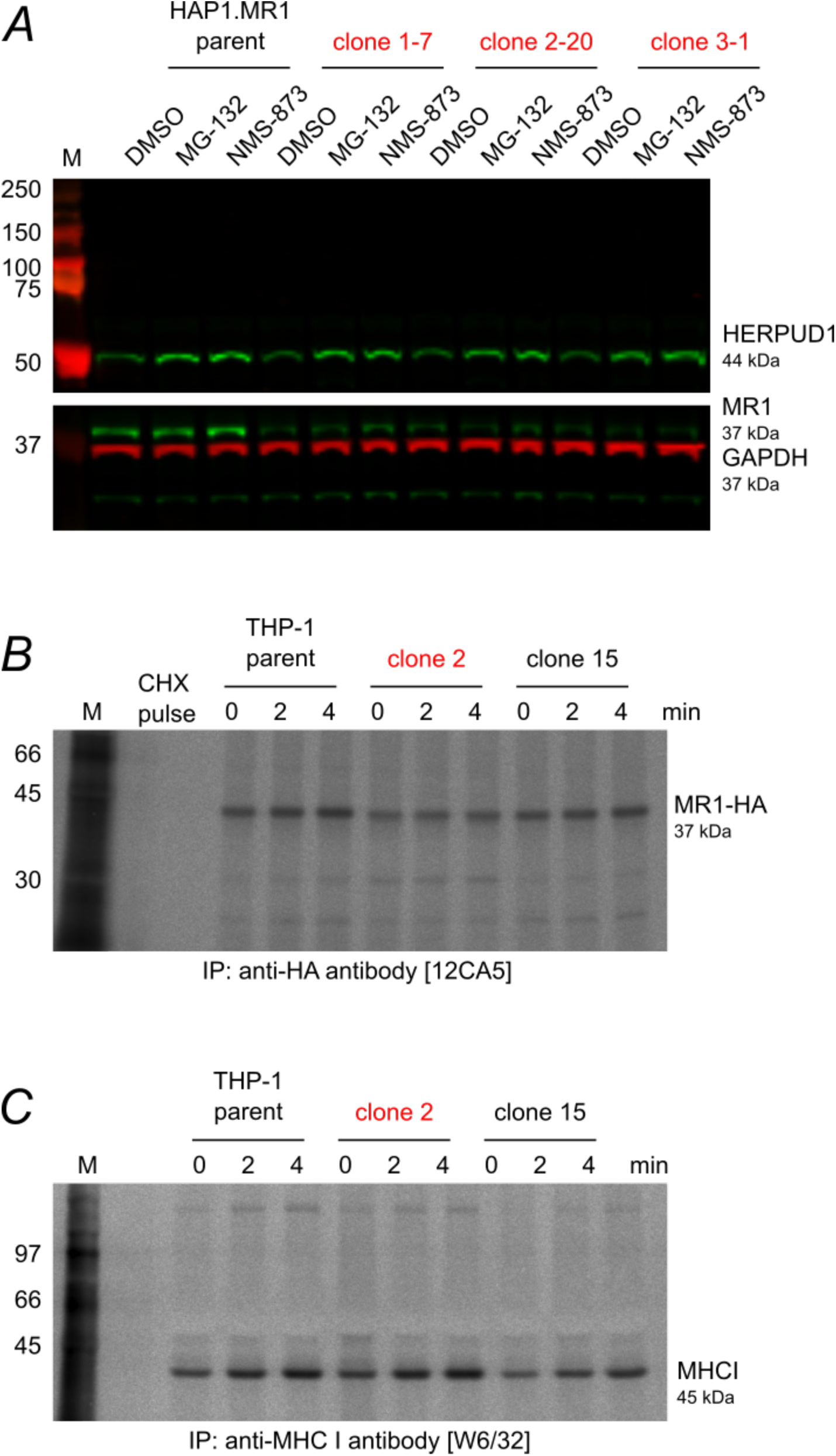
ATP13A1 deficiency affects MR1 levels at or immediately after protein synthesis. *A*, three independent HAP1.MR1 clones and the HAP1.MR1 parent clone D9 (HAP1.MR1 parent) were incubated with MG-132 or NMS-873 for 4 h in duplicates. Protein lysates were separated by SDS-PAGE and immuno-blotted for HERPUD1 (green, top panel), MR1 (green, bottom panel), and GAPDH (red, bottom panel) with the duplicates on two separate gels. Western Blot of one of the two duplicate gels is shown. Membranes were cut prior to primary antibody incubation. M denotes protein marker and molecular weights of visible marker bands are indicated in kDa to the left of the blot. *B* + *C*, THP-1 parent clone, ATP13A1 KO clone 2, and ATP13A1-sufficient clone 15 over-expressing HA-tagged MR1 were pulsed with radioactively labeled cysteine and methionine for 1 min and then chased in the presence of cold Cys/Met and 100 µM cycloheximide for 0, 2, or 4 min before lysis and immuno-precipitation with anti-HA antibody 12CA5 (*B*) and then anti-MHC I antibody W6/32 (*C*). Immuno-precipitated proteins were separated by SDS-PAGE, transferred onto PVDF membranes, and visualised by autoradiography. Cycloheximide was included during the pulse in the control sample (CHX pulse). Molecular weight indications were inferred based on protein marker on the PVDF membranes. ATP13A1 KO clones are highlighted in red. Data in *B* + *C* are representative of two independent experiments.

To further characterize the effect of ATP13A1 on the kinetics of MR1 expression, we performed pulse-chase experiments after metabolic labeling of nascent proteins with radioactive amino acids. In order to be able to immunoprecipitate nascent MR1 molecules regardless of their folding state, we overexpressed human influenza haemagglutinin (HA)-tagged MR1 in one THP-1 ATP13A1 KO clone and one THP-1 ATP13A1-sufficient clone as well as the THP-1 parent clone (Supporting Information Figure 12). Confirming the phenotype seen in the HAP1.MR1 ATP13A1 KO clones, total protein levels of recombinantly overexpressed MR1-HA were lower in the absence of ATP13A1 compared to the THP-1 WT parent clone and the ATP13A1-sufficient clone (Supporting Information Figure 12*B*).

Immuno-precipitation of a radiolabelled cohort of proteins revealed that the levels of MR1-HA were reduced in the ATP13A1 KO cells as early as two minutes after the pulse (Figure 6*B*). Indeed, in the absence of ATP13A1 the HA-reactive population of molecules incorporated the radioactive amino acids less efficiently compared to the parent or the ATP13A1-sufficient cells as evident from the lower signal at time = 0 min (Figure 6*B*). This indicates that ATP13A1 modulates either the rate of MR1 protein synthesis or the stability of the nascent MR1 polypeptide immediately after its co-translational translocation into the ER. Importantly, the abundance and kinetics of folded MHC class I molecules were comparable between all cell lines (Figure 6*C*).

To exclude the possibility that the reduced MR1-HA expression in the ATP13A1 KO background was caused by differences in transduction efficiency, we transduced the clones with an MR1-Emerald construct featuring an internal ribosomal entry site (IRES) (96). In these cells, MR1 and Emerald are co-transcribed but not fused on the protein level. Thus, expression of the green fluorescent protein can be used as an indicator of transduction efficiency without affecting MR1 protein function. Although overall transduction efficiency was indeed slightly lower in the THP-1 ATP13A1 KO clone 2 (Figure 7*A*+*B*), MR1 surface levels were reduced in these cells compared to the THP-1 parent and the ATP13A1-sufficient clone 15 across a range of transduction levels (Figure 7*C*+*E*). Importantly, MHC class I surface expression was not affected (Figure 7*D*+*F*). In conclusion, in the absence of ATP13A1, we observed reduced expression of endogenous MR1 as well as of three exogenously expressed constructs, corroborating the conclusion that MR1 protein levels are stabilized by the P_5_-type ATPase.

**Figure 7:**
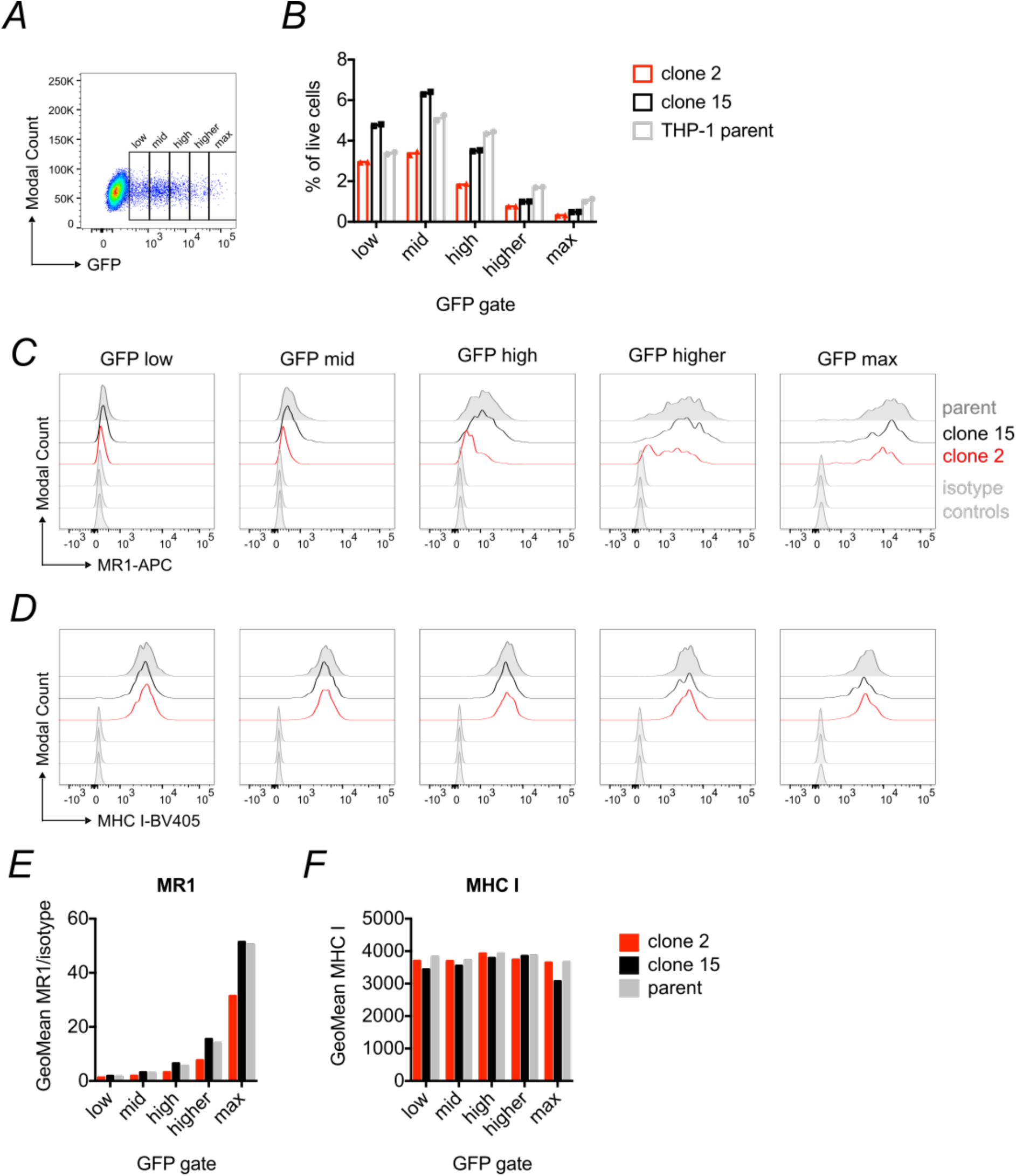
MR1 surface expression is reduced upon loss of ATP13A1 over a range of MR1 expression levels. The THP-1 parent clone, the THP-1 ATP13A1 KO clone 2, and the THP-1 ATP13A1 sufficient clone 15 were transduced with lentiviral particles encoding MR1 and Emerald green fluorescent protein linked by an internal ribosomal entry site (IRES) to compare MR1 surface levels across transduction efficiencies by flow cytometry. *A*, gates capturing different GFP expression levels. *B*, percentage of live cells that fall into each GFP gate in the three cell lines. *C* + *D*, histograms of MR1 (*C*) and MHC class I (*D*) surface levels for each GFP gate and cell line. Geometric means of data shown in *C* + *D* are displayed in *E* + *F*, respectively. Data in *B* are mean of technical duplicates (combined from stained samples and isotype controls). ATP13A1 KO clones are highlighted in red.

## Discussion

MAIT cells are one of the most recent additions to the family of innate-like, unconventional T cells (97). Unlike classical, MHC class I and II-restricted T cells, they are characterised by a limited TCR repertoire, reactivity to conserved, non-peptidic antigens, and an effector-memory phenotype, which allows them to rapidly exert effector functions (15, 22). MAIT cells have been implicated in the control of numerous infectious diseases (recently reviewed in (15)) and emerging data extend their activity to protective as well as pathogenic roles in sterile inflammation, autoimmunity, tissue repair, and cancer (98–104). Their unusually high abundance in both blood and tissues, combined with their restriction by a monomorphic antigen presenting molecule makes them a particularly attractive therapeutic target (24, 25, 28, 103, 104). However, to effectively leverage the potential of these cells in prophylactic and therapeutic applications, a better understanding of the MR1-MAIT cell axis is needed. In particular, the cellular and molecular mechanisms governing MR1 loading and trafficking are incompletely defined (10, 31, 39, 41).

Here, we performed a gene trap screen and identified 199 putative modulators of MR1 surface expression in the presence of the stabilising ligand Ac6FP. The hits included both proteins known to be involved in antigen presentation and protein trafficking and those not previously associated with this process. Intriguingly, classical MHC class I and components of the PLC (TAP1, TAP2, and tapasin) were putative negative regulators of MR1 surface expression in our screen. This differs from a recent CRISPR screen performed by McWilliam, Mak et al., who found tapasin, calreticulin, and TAP among the positive regulators of MR1 surface expression (36). In contrast to earlier data (105), McWilliam, Mak, and colleagues show that MR1 surface expression is reduced in tapasin-deficient cell lines and primary cells. Furthermore, their study confirms previous reports of physical interaction of MR1 and tapasin by co-immunoprecipiation (32, 36). Although shown to physically interact with MR1 in the ER (36), neither calnexin (CANX; log2MI = 0.687; fcpv = 0.442) nor calreticulin (CALR; log2MI = 2.375; fcpv = 0.124) or TAPBPR (TAPBPL; log2MI = 1.159; fcpv = 1) were statistically significantly enriched in our screen for modulators of MR1 surface expression. This discrepancy could be due to technical differences such as using a gene-trap approach rather than the CRISPR/Cas9 system or reflect biological differences between the model systems used. The differing observations emphasize the need for further research in this area.

Due to its localisation in the ER, its predicted transporter function, and its high enrichment in the MR1^low^ population in this study, we focused on the P_5_-ATPase ATP13A1. The data presented here support the conclusion that this transmembrane helix dislocase modulates the size of the intracellular pool of MR1 and consequently, MR1 surface expression and antigen presentation. ATP13A1 is the mammalian orthologue of the yeast ATPase Spf1p (also known as Cod1p) and is unique among the five known mammalian P_5_-ATPases (80). The only member of the P_5_A subgroup in humans, it differs from its P_5_B paralogues ATP13A2-5 in its intron/exon structure and is predicted to have resulted from an earlier gene duplication than the other isoforms (80, 106). In addition, ATP13A1 features a stretch of 67 amino acids which is not present in the other mammalian isoforms in either human or mouse but in the yeast, *Caenorhabditis elegans*, and *Drosophila melanogaster* orthologues (79, 80). Like all P-type ATPases, ATP13A1 features a central DKTGTLT motif containing the catalytic aspartate (77, 79, 80, 107) and it also contains conserved nucleotide binding domains (79, 80). Topology predictions have identified at least 10 putative transmembrane domains for mammalian ATP13A1 (80). Both ATP13A1 and ATP13A4 localise to the ER and have been reported to be involved in calcium homeostasis (69, 74, 108). However, direct calcium transport activity has not been shown for ATP13A1 and ATPase activity of purified Spf1p was not dependent on or enhanced in the presence of calcium in vitro (74). The concentration of manganese in microsomes isolated from Δ*spf1* cells was decreased compared to that in wild type microsomes, though other divalent cations such as calcium and magnesium were not tested in this study (69).

In addition to their implication in ER ion homeostasis, P_5_A-ATPases are associated with a wide variety of cellular functions including ER stress and the UPR, protein processing and turnover, membrane insertion and targeting of secretory proteins, intracellular sterol distribution and vesicular transport, and many more (109). Despite, or perhaps because of, these pleiotropic phenotypes, the substrate of this group of transporters remained enigmatic until very recently (109). In a landmark publication, McKenna et al. demonstrated that human and yeast P_5_A-ATPases act as protein dislocases, removing mis-targeted tail-anchored mitochondrial proteins wrongly inserted into the ER membrane (73). In this manuscript we show that loss of ATP13A1 corresponded to a drastic decrease in total cellular MR1 levels in two different cell lines and examined possible mechanisms by which this transporter may impact the amount of MR1 available in the cell.

Since *spf1* mutants suffer severe ER stress (69, 74, 76–78), we hypothesized that the MR1 phenotype we observed in the ATP13A1-deficient cells may be caused by ER-associated degradation due to misfolding of either MR1 itself or a crucial chaperone. However, we did not find any evidence for elevated ER stress in ATP13A1-deficient HAP1.MR1 cells and inhibition of proteasomal degradation did not rescue MR1 levels in these cells at steady state. This indicates that MR1 molecules do not accumulate in an unfolded state in the ER, which would necessitate clearance by the UPR and is in stark contrast to the yeast system (74, 77, 78, 110). Of note, the metazoan ER stress response comprises three overlapping branches while the equivalent system in yeast cells relies on IRE1 sensing only (111). With less in-built redundancy, this simpler system may be less robust, providing a potential explanation as to why yeast cells appear to be more dependent on Spf1p for maintaining ER homeostasis. Alternatively, ER stress may be a downstream effect of Spf1p deficiency rather than a direct consequence of it and mammalian cells may have more sophisticated mechanisms to counteract such secondary effects.

Early investigations identified a hypoglycosylation phenotype in *spf1* mutants (69, 75, 76). Yet, the small population of MR1 molecules detectable in HAP1.MR1 ATP13A1 KO clones was still sensitive to endoglycosidase digestion, confirming that glycoprotein processing in the ER was not affected upon loss of the transporter. This observation is consistent with reports that EndoH-sensitive core glycosylation is intact in *spf1* KO yeast strains and only higher order polysaccharide residues could not be processed in these mutants (76).

Since the reduction of total MR1 protein was not accompanied by a statistically significant difference on the mRNA level, we focussed our efforts on post-transcriptional events. Pulse-chase experiments revealed that the difference in total MR1 protein levels was detectable within minutes of translation and affected MR1 molecules regardless of their folding state. This is consistent with a model where ATP13A1 function is important for the stabilisation of nascent MR1 polypeptides immediately after translation and suggests a very early bifurcation into a cohort of MR1 molecules that successfully complete protein synthesis and initial folding and a cohort of molecules that is immediately degraded. Alternatively, the rate of MR1 protein synthesis may be affected in these cells. The observed trend that radioactive labelling of MR1 was less efficient in an ATP13A1 KO clone at early time points compared to THP-1 WT cells or an ATP13A1-sufficient clone derived from the same parent supports this hypothesis. Intriguingly, protein translocation into the ER through Sec61 is impaired in an *spf1* yeast mutant (112), providing a potential mechanistic basis for this model. However, such an effect on protein translocation would be expected to have a global effect, yet antigen presentation by classical MHC class I and CD1d was unaffected in our ATP13A1 KO cells. Pertinent to this, a recent siRNA screen to identify modulators of CD1d-mediated antigen presentation did not find ATP13A1 (84). A recent CRISPR screen for MR1 chaperones, however, also found sgRNAs targeting ATP13A1 to be enriched in cells with low MR1 surface expression (36), further supporting the role for this ATPase described here.

Although Spf1p has been shown to modulate ion concentrations in the ER, it has also been postulated that its substrates may be more diverse, potentially even including macromolecules such as aminophopholipids based on sequence similarities to P_4_-type ATPases (74, 109). Since it is still unclear how soluble MR1 ligands access the ER where they encounter partially folded, ligand-receptive MR1 molecules (37, 41) it is tempting to speculate that ATP13A1 may also transport vitamin metabolites, implying a role analogous to TAP which transports antigenic peptides into the ER for loading onto MHC class I molecules (113). Pertinent to this, Sørensen and colleagues recently identified phopsphatidyl-inositol 4-phosphate as a potent stimulator of Spf1p ATPase activity and hypothesised that sterol flippase activity of P_5_A-ATPases may underlie the diverse phenotypes observed upon mutation of the transporters (112). Interestingly, two putative negative regulators of MR1 surface expression identified in our screen (VAC14; log2MI = 2.087; fcpv = 7.39×10^-6^; and FIG4; log2MI = 1.879; fcpv = 1.05×10^-3^) are components of a complex that regulates the interconversion of phosphatidylinositol-derived signalling lipids (52), further strengthening a potential link between MR1-mediated antigen presentation and sterol homeostasis.

Recent work by McKenna et al. further extended the list of P_5_A-ATPase substrates to short transmembrane helices of mis-localised proteins (73). In this study, the authors obtained the first structural information on Spf1p, which provides an explanation for the unusual substrate selectivity of P_5_A-ATPases: compared to other P-type transporters, Spf1p features an unusually large binding pocket with a lateral opening facing the lipid phase of the ER membrane. This topology is ideally suited for its interaction with short, helical transmembrane segments which the ATPase removes from the lipid bilayer. In this way, tail-anchored mitochondrial proteins that are incorrectly inserted into the ER membrane can be removed and released into the cytosol for re-targeting to the correct organelle. Using quantitative proteomics, the authors show that loss of ATP13A1 leads to decreased protein abundance of a variety of membrane-anchored proteins (73), consistent with our findings on reduced MR1 protein levels in ATP13A1 KO cells. Unfortunately, the proteomics analysis does not include data on MR1 (73). Of note, HLA-A protein levels were not affected (73), consistent with our results. Although mis-oriented N-terminal signal sequences can also be substrates for ATP13A1 (73) and MR1 features such a peptide at is N-terminus, ATP13A1 did not coimmunoprecipitate with MR1 in our hands (data not shown). Similarly, a recent coimmunoprecipitation screen provided no evidence that MR1 directly interacts with the transporter (36).

It remains a likely possibility that the defect in MR1 antigen presentation is an indirect rather than a direct consequence of loss of ATP13A1. Since inhibition of proteasomal degradation did not rescue the MR1 phenotype, it further remains to be determined whether MR1 molecules in the ATP13A1 KO cells are degraded by a different means or whether the rate MR1 protein synthesis is reduced in these mutants. If protein synthesis was affected, further questions would arise as to the mechanism by which the ER transporter regulates translation and, most importantly, how this is specific to MR1.

In summary, we provide evidence that the P_5_-ATPase ATP13A1 is an important cellular factor impacting MR1 surface expression and antigen presentation through a role in MR1 biogenesis in the ER. Our findings suggest that the mechanism underlying this effect does not involve differential glycosylation or general ER stress but appears to act on nascent MR1 polypeptides upon or immediately after protein synthesis. Further investigations into the transcriptional and translational effects of ATP13A1 deficiency as well as the extent to which these effects can be generalised to *in vivo* models are needed to confirm our data and determine the molecular mechanism responsible for the MR1 phenotype.

## Experimental Procedures

### Cell culture

THP-1, C1R, and HEK293T cells were purchased from ATCC. The generation of the Melan A-reactive T cell line was described previously (114). HAP1 WT 1524 was obtained from the Nijman lab (Nuffield Department of Medicine, University of Oxford, UK). MAIT cells and iNKT cells were sorted from human peripheral blood mononucleocytes (PBMCs) isolated from leukocyte cones obtained from the NHS Blood and Transplant Unit as described below. This work was covered by HTA licence number 12433. Cells were kept at 37° C with 5% CO_2_. In general, THP-1 cells and their derivatives were cultured in RPMI-1640 (Sigma-Aldrich or Gibco) supplemented with 10% Foetal calf serum (FCS; Sigma-Aldrich or Gibco), 2 mM Glutamine (Sigma-Aldrich), 1 mM Sodium Pyruvate (Gibco), 1x Non-essential amino acids (Gibco), 100 U/ml penicillin + 100 µg/ml streptomycin (Sigma-Aldrich), 10 mM HEPES (Gibco), and 50 µM β-mercaptoethanol (Gibco). HEK293T cells were cultured in DMEM (Sigma-Aldrich) supplemented in the same way. Sorted human MAIT cells and T cells were cultured in IMDM (Gibco) supplemented with 5% pooled human serum (NHS Blood and Transplant Unit), approximately 5000 U/ml IL-2 (as described (115)), and supplements as above. HAP1 cells and their derivatives were cultured in IMDM (Gibco) supplemented with 10% FCS and 100 U/ml penicillin + 100 µg/ml streptomycin only. Cells were sub-cultured every 2-4 days depending on confluency. Adherent cells were detached using either PBS + 1 mM EDTA (VWR) or Trypsin-EDTA (Gibco). Cells were stored in 10% DMSO (Sigma-Aldrich) in FCS in liquid nitrogen storage tanks for long term storage.

### PBMC isolation and MAIT/iNKT cell sorting

PBMCs were isolated from leukocyte cones obtained from the NHS Blood and Transplant Unit. Samples with suitable MAIT cell numbers as determined by flow cytometry were enriched for CD2^+^ cells by MACS sorting (Miltenyi) according the manufacturer’s protocol. A representative enriched sample was >95% positive for CD2 as determined by flow cytometry (eBioscience, clone RPA-2.10). Cells were stored at 4° C o/n before surface staining for CD3 (eBioscience, clone SK7; not always included), CD161 (eBioscience, clone HP-3G10) and the Vα7.2 TCR (BioLegend, clone 3C10) to enrich for Vα7.2^+^ CD161^high^ MAIT cells on a SONY MA900, a FACSAria Fusion, or a FACSAria III sorter (both BD Biosciences) in the WIMM Flow Cytometry Facility. Sort purity was routinely >90%. Invariant natural killer T (iNKT) cells were enriched with the antibody clone 6B11 (BioLegend) and maintained as described (116).

### Generation of over-expressing cell lines

To generate cell lines over-expressing human MR1, the pHR-SIN-MR1-IRES-Emerald lentiviral expression vector previously made in our lab (96) was digested with XhoI and NotI endonucleases and the Emerald coding sequence was replaced with a 21 base pair adapter designed to include a BamHI restriction site to enable screening by PCR. The undigested plasmid was used for the IRES-Emerald transductions. The plasmid encoding the HA-tagged MR1 has been described previously (5). Briefly, the MR1 coding region was amplified from the pHR-SIN-MR1-IRES-Emerald vector (96) with a forward primer having a BglII overhang and a reverse primer introducing an HA sequence and a NotI restriction site. This was then cloned into the pHR-SIN-IRES-Emerald vector using the BamHI and NotI restriction sites upstream and downstream of the IRES-Emerald cassette, respectively. This vector was derived from the pHR-SIN-CSGW plasmid (117, 118). Of note, the MR1 coding sequence used carries two silent mutations: A138G and C939T. All oligonucleotide sequences can be found in Supporting Information Table 4. The lentiviral expression vector for human ATP13A1 was purchased from VectorBuilder as a bacterial stock (pLV[Exp]-Bsd-{sffv}>hATP13A1[NM_020410.2]). Lentiviral particles were produced in HEK293T cells using packaging vectors encoding gag/pol (pCMVR8.91) and vsv-g (pMDG). Polyclonal MR1-overexpressing cell lines were flow sorted based on MR1 surface expression. ATP13A1 over-expressing cells were selected with Blasticidin S (Sigma-Aldrich).

### Flow Cytometry

Cells were incubated with Acetyl-6-formylpterin (Schirks Laboratories) as specified in the figure legends, harvested, and transferred to round-bottom 96-well plates for staining with Aqua live/dead stain (Life Technology or Biolegend), followed by Fc receptor blocking (normal immunoglobulin, octopharma or Fc Receptor Binding Inhibitor Antibody, eBioscience), and surface staining on ice. If applicable, cells were split into two wells and primary antibody mix containing either the target antibody or the isotype control antibody was added. When applicable cells were fixed in 2% PFA (Electron Microscopy Sciences) or ICS fixation buffer (eBioscience) before acquisition. Buffer containing 2 mM EDTA was used throughout the procedure for HAP1 cell lines. On occasion, Propidium Iodide (Sigma-Aldrich) was added to samples just before acquisition instead of staining with aqua live/dead stain. For ploidy staining, cells were fixed in 70% ethanol and stained with 10 µg/ml Propidium Iodide (PI) in the presence of 0.5 mg/ml RNase A before analysis on a flow cytometer in the 561-610/20 or the 561-620/15 channel on linear scale. Single colour controls were either cells or OneComp eBeads Compensation Beads (Life Technologies). Cells were analysed on an Attune NxT analyser equipped with four lasers (Life Technologies), a CyAn ADP analyser (Beckman Coulter) or an LSR Fortessa X50 fitted with five lasers (BD Biosciences) in the WIMM Flow Cytometry Facility and data were analysed and displayed using FlowJo v9 or v10 (FlowJo). Y-axes of histograms show modal counts and x-axes are in logarithmic or biexponential scale except for FSC and SSC which are on a linear scale. Antibodies used were clone 26.5 for MR1 (BioLegend), clone MOPC-173 as IgG2A control (BioLegend), and clones W6/32 (BioLegend) and G46-2.6 (BD Biosciences) for MHC class I. Fluorophores are indicated on the plot axes.

For analysis of HAP1 CRISPR lines generated for bulk validation, cells were dissociated with 0.05% Trypsin-EDTA (Thermo Fisher Scientific), counted using a TC20 automated cell counter (Bio-Rad) and cell number across samples was equalized. Cells were incubated with primary antibody dilution for 1 hour at 4°C. After 2 washes, cells were incubated with Alexa488-conjugated secondary antibody for 45 minutes at 4°C. Cells were fixed in ICS Fixation Buffer for 30 minutes at RT, washed twice, and incubated in 1X Permeabilisation Buffer (BD Bioscience) for 10 min at RT. Cell pellet was then washed and incubated with 0.8 mg/ml RNase A (Applichem) for 1.5 hours before 2 additional washing steps and incubation with 10 µg/ml PI. According to the manufacturer instruction, RNAse A and PI incubation steps were performed in 1X Permeabilisation Buffer.

### Antigen presentation assays

For functional assays, antigen presenting cells (APCs) were incubated with the indicated stimuli for 5 h at 37° C. Technical duplicates were split either before or after pulsing with the antigen such that the final number of APCs was 50 000 per well. For stimulation with *E.coli*, DH5α bacteria were grown in Luria Broth over night, washed in PBS, and diluted to an OD_600_ of 0.5, corresponding to approximately 4×10^8^/ml as calculated based on the online Agilent Biocalculator (119). Stimulation with live bacteria was carried out in medium without antibiotics and gentamycin (Lonza) was added to a final concentration of 50 µg/ml after 3 h. Peptide (ELAGIGILTV, purchased from Sigma-Aldrich) pulsing was performed in serum-free medium. α-Galactosylceramide (αGalCer) was purchased from Enzo Life Sciences. 5-A-RU was synthesised as described previously (120) and combined with methyl-glyoxal (Sigma-Aldrich) immediately before addition to the culture. Stimuli were washed off before co-incubation with either 15 000 or 20 000 T cells per well for at least 36 h. Interferon γ (IFNγ) in the culture supernatants was measured by ELISA using commercially available antibody pairs (BD Biosciences, clones NIB42 for the capture antibody and 4S.B3 for the biotinylated detection antibody). Antigen binding was visualized with avidin-peroxidase (Sigma-Aldrich) and developing solution containing phenylenediamine (Sigma-Aldrich). The reaction was stopped with 2 M H_2_SO_4_ (Fluka) when the standard curve had fully developed. Colour intensity was measured at 490 nm in an iMark plate reader (Bio-Rad) and quantified using a 4-parameter standard curve. Supernatants were diluted as appropriate and added in a final volume of 25 µl/well. Standard curves of recombinant IFNγ (PeproTech) were in 50 µl/well.

### Western blots

HAP1 cells were harvested for Western blots with Trypsin/EDTA or PBS + 1 mM EDTA to preserve the surface proteome. Sometimes cells were lysed by adding lysis buffer directly to the dish. To make protein lysates cells were washed in PBS, pelleted, and lysed at 4° C in either RIPA buffer (Sigma-Aldrich) or self-made lysis buffers (either 50 mM Tris pH 7.5, 150 mM NaCl, 1 mM EDTA, 0.5% NP40 (IGEPAL CA-630) or 20mM Tris-HCl pH 7.5, 150mM NaCl, 1mM EDTA pH 8.0, 1% Triton X-100, 1mM EGTA, 2.5mM sodium pyrophosphate). All lysis buffers were supplemented with protease inhibitors (Roche). Cell lysates were cleared by centrifugation and protein-containing supernatants were collected into fresh tubes and quantified with the Pierce BCA protein assay kit (Thermo Scientific) using a two-fold dilution curve of BSA (Thermo Scientific) as a standard. HAP1 WT or D9 parent control lysates were not always prepared on the same day as the clone lysates. Samples were prepared in 1x Loading Buffer and 1x Reducing Agent (both Invitrogen) to contain equal amounts of protein, boiled, and loaded onto a gel according to their size. Generally, proteins with a molecular weight (MW) below 100 kDa were separated on 4-12% Bis/Tris gels (Life Technologies) in MES SDS Buffer (Invitrogen) and proteins with a molecular weight above 100 kDa were separated on 3-8% Tris/Acetate gels (Life Technologies) in MOPS Buffer (Invitrogen). Appropriate MW markers were included. Gels were transferred onto a PVDF or nitrocellulose membrane (both Bio-Rad) using the TransBlot Turbo Transfer system (Bio-Rad), and blocked in PBS + 0.1% Tween-20 + 5% bovine serum albumin (BSA) (Sigma-Aldrich or Fisher Scientific). If needed, membranes were cut to size for incubation with different primary antibodies. Membranes were washed in PBS + 0.1 or 0.5% Tween-20 (Acros Organics), incubated with LI-COR® secondary antibodies with agitation, washed, and dried between tissues before imaging with the Odyssey® Near-Infrared imaging system. For Endoglycosidase H (EndoH, NEB) digestions, equal amounts of protein from RIPA cell lysates were denatured in supplied Denaturation Buffer before splitting each sample in two. One half was incubated with the enzyme in the appropriate Glycobuffer and the other half with Glycobuffer only. Loading buffer and reducing agent were added to each reaction and samples were analysed by SDS-PAGE as above. Primary antibodies were clone 6C5 (mouse, Santa Cruz) or D16H11 (rabbit, Cell Signaling Technology) for GAPDH, C4 (Santa Cruz) for ACTB, polyclonal (Cat. No. 13260-1-AP; rabbit, Proteintech) for MR1, polyclonal (Cat. No. 16244-1-AP; rabbit, Proteintech) for ATP13A1, clone HC10 (mouse, purified from hybridoma supernatant by HIU lab management) for MHC class I, and clone EP2978Y (rabbit, Abcam) for β_2_m. Suitable IRDye 680RD and 800CW secondary antibodies were purchased from LI-COR.

For analysis of HAP1 cells for bulk validation, cells were lysed and collected in RIPA buffer (Sigma-Aldrich) supplemented with protease and phosphatase inhibitors (Roche, Sigma-Aldrich). Lysates were mixed 1:1 with 2x reducing sample buffer (320 mM Trizma base (pH adjusted to 6.8), 40% glycerol, 16 µg/ml bromophenol blue, 8% SDS) and boiled for 10 minutes. Proteins were separated on 4-12% Bis-Tris Gels (Invitrogen) and transferred to PVDF membrane (Biorad). A solution of 0.2% I-Block (Applied Biosystems) in PBS + 0.1% Tween-20 (Sigma-Aldrich) (PBST) was used for blocking and antibody incubation, washing was performed in PBST. Species-specific HRP-conjugated antibody (BioRad) signal was detected using Western Lightning Plus-ECL (Perkin Elmer) and imaged on a Universal Hood III machine (Bio-Rad).

### qRT-PCR

RNA was extracted using either the QIAGEN RNeasy kit or the Ambion RNAqueous kit according to the manufacturer’s instructions including the respective DNase digestion protocol. RNA was quantified by Nanodrop (Thermo Scientific) and equal amounts of RNA were reverse transcribed using Ambion or Takara Bio kits. Only cDNA transcribed with the same kit was directly compared in qRT-PCR reactions. HAP1 WT or D9 parent control RNA was not always extracted on the same day as the clone RNA. Pooled “no RT” and “no RNA” controls were routinely included and occasionally gave a signal but always at a much higher C_T_ value than the samples. qRT-PCRs were run as technical duplicates or triplicates on a QuantStudio7 qRT-PCR machine (Life Technologies) and expression was normalised to a house keeping gene (HKG). Taqman probes used can be found in Supporting Information Table 4.

### ER stress assay

HAP1.MR1-derived clones were treated with 0.1 µM thapsigargin (Sigma-Aldrich) or solvent control DMSO for 3 h. Protein lysates were obtained and Western Blots performed as above. In the same experiments, RNA was extracted and reverse transcribed as above. XBP1 cDNA was PCR amplified with previously published primers (121) (Supporting Information Table 4) and analysed for the XBP1 splice variant on a 2.5% agarose gel.

### Metabolic labelling and Pulse-Chase experiments

For metabolic labelling, 1.5 x 10^6^ THP-1 cells were starved in the absence of methionine and cysteine for 1 h. Following starvation, cells were pulse labelled with 50µCi of ^35^S-methionine and -cysteine for 1 min (EasyTag™ EXPRESS35S Protein Labeling Mix, Perkin Elmer, Waltham MA, USA) and immediately transferred into 10x volume of chase medium containing excess amounts of unlabelled cysteine (500 µg/ml), methionine (100 µg/ml) and 100 µM cycloheximide to prevent incorporation of radioactive amino acids. Cells were harvested at 0, 2 and 4 minutes and lysed in native lysis buffer (50 mM Tris·Cl, pH 7.5,150 mM NaCl, 5 mM EDTA, 1% (v/v) Triton X-100) for 1 h. Hemagglutinin-tagged MR1 was immunoprecipitated using monoclonal anti-HA antiserum (clone 12CA5). HLA class I proteins were immunoprecipitated from the supernatant of the first immunoprecipitation with anti-HA, using W6/32 antibody.

### Gene trap mutagenesis and sort

Gene trap mutagenesis was carried out as described before (44) with small changes. Briefly, HEK293T cells were co-transfected with a gene trap vector encoding for a strong adenoviral splice acceptor as well as blue fluorescent protein (pGT-en2-BFP344) and retroviral packaging plasmids Gag-pol, VSVg, and pAdv (43). Virus-containing supernatant was harvested and concentrated by ultracentrifugation twice a day for three consecutive days. Concentrates from each day were stored at 4° C o/n and then pooled for transfection. 2×10^7^ cells of HAP1.MR1 clone D9 were seeded in a T175 flask on Day 0 and transduced with concentrated gene-trap retrovirus in the presence of 8 µg/ml of protamine sulphate (MP Biomedicals #P4380) for three consecutive rounds of transduction on Days 1-3 before expansion and freezing. Expanded transduced HAP1.MR1 D9 cells were thawed and approximately 3×10^9^ cells were pulsed with 5 µg/ml Ac6FP for 5 h before cell surface staining with anti-MR1 primary antibody (clone 26.5) and goat anti-mouse secondary antibody coupled to Alexa488 (Life Technologies). Stained cells were fixed, permeabilised, and stained with Propidium Iodide before sorting on a LE-SH800 sorter (Sony) fitted with a 100 µm chip over four days gating on haploid cells based on PI staining. MR1^hi^ and MR1^low^ populations comprising approximately 1% of total, respectively, were sorted (Supporting Information Figure 2).

### Recovery of insertion sites and bioinformatics analysis

Gene trap insertion sites were recovered using linear amplification-mediated polymerase chain reaction (LAM-PCR) similarly to the protocol described by Blomen et al. (122) and Brockmann et al. (44). Briefly, sorted MR1^hi^ and MR1^low^ cells were de-crosslinked and DNA was extracted from the two populations separately. Long terminal repeat (LTR)-proximal gene regions were amplified from 4×1 µg genomic DNA in four separate reactions per MR1 population using a double-biotinylated primer. Biotinylated single-stranded DNA (ssDNA) was isolated and Illumina sequencing adapters were added. Libraries were PCR-purified and sequenced on a HiSeq2500 machine (Illumina) generating 50 base pair (bp) single end reads with the 5’ sequencing primer. All primer sequences can be found in Supporting Information Table 4. Recovered sequences were analysed with the pipeline described by Brockmann et al. (44). In short, reads were mapped onto the human reference genome hg19 by running the aligner Bowtie (123) twice, allowing for 0 and 1 bp mismatches to avoid omitting reads that do not map uniquely when tolerating 1 bp mismatch. Reads were mapped using hg19 protein-coding gene coordinates (Refseq) and intersectBED (124). A customised BED file was used for both gene mapping and determining the orientation of the integration relative to the gene (sense or antisense). Only sense insertions in non-overlapping gene regions outside the 3’ untranslated region (UTR) were considered to have the potential to disrupt gene function and taken into account for further analysis. The number of unique disruptive insertions for a gene in the MR1^hi^ or MR1^low^ population was normalised to the total number of insertions in the respective populations and compared to the number of normalised insertions for that gene in the opposite population using a two-sided Fisher’s exact test. Significance of enrichment in either of the two tails was determined by applying the Benjamini–Hochberg false discovery rate correction to the calculated p-values. In addition, enrichment in either tail was expressed as the Mutation Index (MI), calculated for each gene as shown below. If a gene was sequenced in only one of the populations, one insertion was added in order to enable calculation of the mutational index.

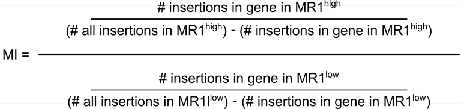

Since functional MR1 protein was only produced from the exogenous MR1 cassette, calculating the MI based on insertions in the entire gene body (log2MI = −1.162, fcpv =7.41×10^-8^) would underestimate enrichment in the MR1^low^ fraction. Thus, the MI for MR1 was re-calculated considering only insertions in exons (log2MI = −2.204, fcpv = 2.90×10^-14^).

### Generation of ATP13A1 KO clones

For bulk validation in HAP1.MR1 parent clone D9, knockout cell lines were generated using the Clustered Regularly Interspaced Short Palindromic Repeats/ CRISPR associated protein 9 (CRISPR/ Cas9) system. LentiCRISPR v2 plasmids encoding for *Streptococcus pyogenes* Cas9 and gene-specific single guide RNA (sgRNA) were generated according to the protocol in (PMID: 25075903 and PMID: 24336571). Target sequences were identified based on the Sabatini library(125), TKO v3 (http://tko.ccbr.utoronto.ca)(126), and GenomeCRISPR (http://genomecrispr.dkfz.de). HAP1 cells were transfected with 2 µg lentiCRISPR v2 bearing the sgRNA of interest in the presence of transfection reagent Turbofectin 8.0 (OriGene). Transfected cells were selected with 2 µg/ml puromycin (InvivoGen) until death of non-transfected control.

For in-depth validation of ATP13A1, three one-component CRISPR/Cas9 plasmids with a different single guide RNA (sgRNA) target sequence within ATP13A1 each were designed based on the Sabatini/Lander CRISPR pooled library (125) (Supporting Information Table 4, Supporting Information Figure 4). sgRNA sequences were tested for specificity using the UCSC-genome browser BLAT tool (https://genome.ucsc.edu/cgi-bin/hgBlat) and cloned into the pX458_Ruby backbone (Addgene #110164). HAP1.MR1 parent clones or a THP-1 WT clone obtained by limiting dilution were transfected with CRISPR guides and sorted based on mRuby expression after 2-3 days. Single cells were sorted into 96-well plates and clones were picked for expansion approximately two weeks after sorting. For sequencing of CRISPR clones, DNA was extracted using the QIAGEN Blood and Tissue Kit in combination with Ambion Proteinase K. For near-haploid HAP1 clones, genomic DNA around the sgRNA target site was amplified and amplicons were sent for Sanger sequencing at the HIU Sequencing Facility and sequences were analysed with SnapGene Viewer (GSL Biotech) and SerialCloner (SerialBasics). For polyploid THP-1 ATP13A1 KO clones, regions around the sgRNA target sited were amplified from genomic DNA using primers that generated overhangs compatible with Illumina adapters (Supporting Information Table 4). PCR products were gel-purified (QIAGEN) and bar-coded using the Illumina index adapter series D5 and D7. Barcoded amplicons were analysed on an Agilent 2200 TapeStation System, pooled in equimolar ratios, cleaned up with Agincourt Ampure XP beads (Beckman Coulter), and quantified on a Qubit3 Fluorometer. The library was sequenced paired-end (read distribution: 151 bp each side) on an Illumina Miseq machine using the Miseq Reagent Nano Kit V2 300-cycle (Illumina). Sequences were analysed using the CRISPResso (127) web implementation.

### Data Analysis and Plotting

Data were analysed in Excel for Mac 2011 (Microsoft) and plotted using Prism (GraphPad Software) Version 7 and 8. Statistical analysis was performed using Prism (GraphPad Software) Version 7 and 8. Figures were assembled in Inkscape (inkscape.org).

## Supporting information

SupportingInformation

## Data Availability

Data of repeat experiments is included in the Supporting Information whenever practical. Data not included are available upon request.

## Supporting Information

This article contains supporting information. The SI references Wang *et al*. (125) and Hollien *et al*. (121).

## Acknowledgements

We gratefully acknowledge Kevin Clark, Sally-Ann Clark, Paul Sopp, and Craig Waugh in the flow cytometry facility at the MRC WIMM for providing cell sorting services, members of the Brummelkamp laboratory (Netherlands Cancer Institute, Amsterdam, The Netherlands) for bioinformatics analysis of the gene trap screen, the MRC WIMM Genome Engineering Services for the design and cloning of the CRISPR constructs, Tim Rostron and John Frankland in the Sequencing facility at the MRC WIMM for providing sequencing services, and David Lewinsohn for critical reading of the manuscript. We dedicate this study to the memory of the late Professor V.C., FRS, an inspirational colleague, mentor, and pioneer in the field of iNKT/CD1/MAIT cell biology, who passed away last year.

## Funding and additional information

C.K. was supported by a Wellcome Studentship (105401/Z/14/Z). M.S., C.G-L., and V.C. are supported by MRC Weatherall Institute of Molecular Medicine Human Immunology Unit core funding. V.C. received a CRUK program grant (C399/A2291). J.C.C. is supported by a CRUK Senior Fellowship (C68569/A29217). G.S.B. is supported by MRC grants MR/S000542/1 and MR/R001154/1. P.K. is supported by Wellcome (WT109965MA). S.S. receives funding from the Deutsche Forschungsgemeinschaft DFG (SP583/7-2). The Flow Cytometry Facility at the MRC WIMM is supported by the MRC HIU; MRC MHU (MC_UU_12009); NIHR Oxford BRC; Kay Kendall Leukaemia Fund (KKL1057), John Fell Fund (131/030 and 101/517), the EPA fund (CF182 and CF170) and by the MRC WIMM Strategic Alliance awards G0902418 and MC_UU_12025. The MRC WIMM Genome Engineering Service is supported by grants MC_UU 12009, the John Fell Fund 123/737 and by the WIMM Strategic Alliance awards G0902418 and MC_UU_12025. The MRC WIMM Sequencing facility is supported by the MRC HIU and by the EPA fund (CF268).

## Abbreviations

2ry: Secondary antibody
5-A-RU: 5-amino-ribityl uracil
5-OP-RU: 5-(2-oxopropylideneamino)-6-D-ribitylaminouracil
Ac6FP: acetyl-6-formylpterin
AF488: AlexaFluor-488
αGalCer: α-Galactosylceramide
APC: antigen presenting cell
APC: allophycocyanin
β2m: β2-microglobulin
bp: basepair
BV405: Brilliant Violet 405
cDNA: complementary DNA
CHX: cycloheximide
E. *coli*: Escherichia coli
EndoH: endoglycosidase H
ER: endoplasmic reticulum
ERAD: ER-associated degradation
fcpv: FDR-corrected p value
FCS: Foetal calf serum
FDR: false discovery rate
FMO: fluorescence minus one
FSC: forward scatter
GeoMean: geometric mean fluorescence intensity
HA: human influenza haemagglutinin
HKG: housekeeping gene
HLA: human leukocyte antigen
IFNγ: interferon-γ
IL: interleukin
iNKT: invariant natural killer T cell
IRES: internal ribosomal entry site
KO: knock-out
LAM-PCR: linear amplification-mediated polymerase chain reaction
L/D: live/dead stain
LTR: long terminal repeat
M: molecular weight marker
MAIT: mucosal-associated invariant T cell
MG: methylglyoxal
MHC: major histocompatibility complex
MI: mutation index
MR1: MHC I-related protein 1
MR1T: MR1-restricted T cell
MW: molecular weight
noRT: no reverse transcriptase control
PBMC: peripheral blood mononucleocytes
PI: Propidium Iodide
PFA: Paraformaldehyde
PLC: peptide loading complex
sgRNA: single guide RNA
SSC: side scatter
ssDNA: single stranded DNA
TCR: T cell receptor
TG: thapsigargin
UPR: unfolded protein response
UTR: untranslated region

