## SupportingInformation for "The P_5_-ATPase ATP13A1 modulates MR1-mediated antigen presentation"

for

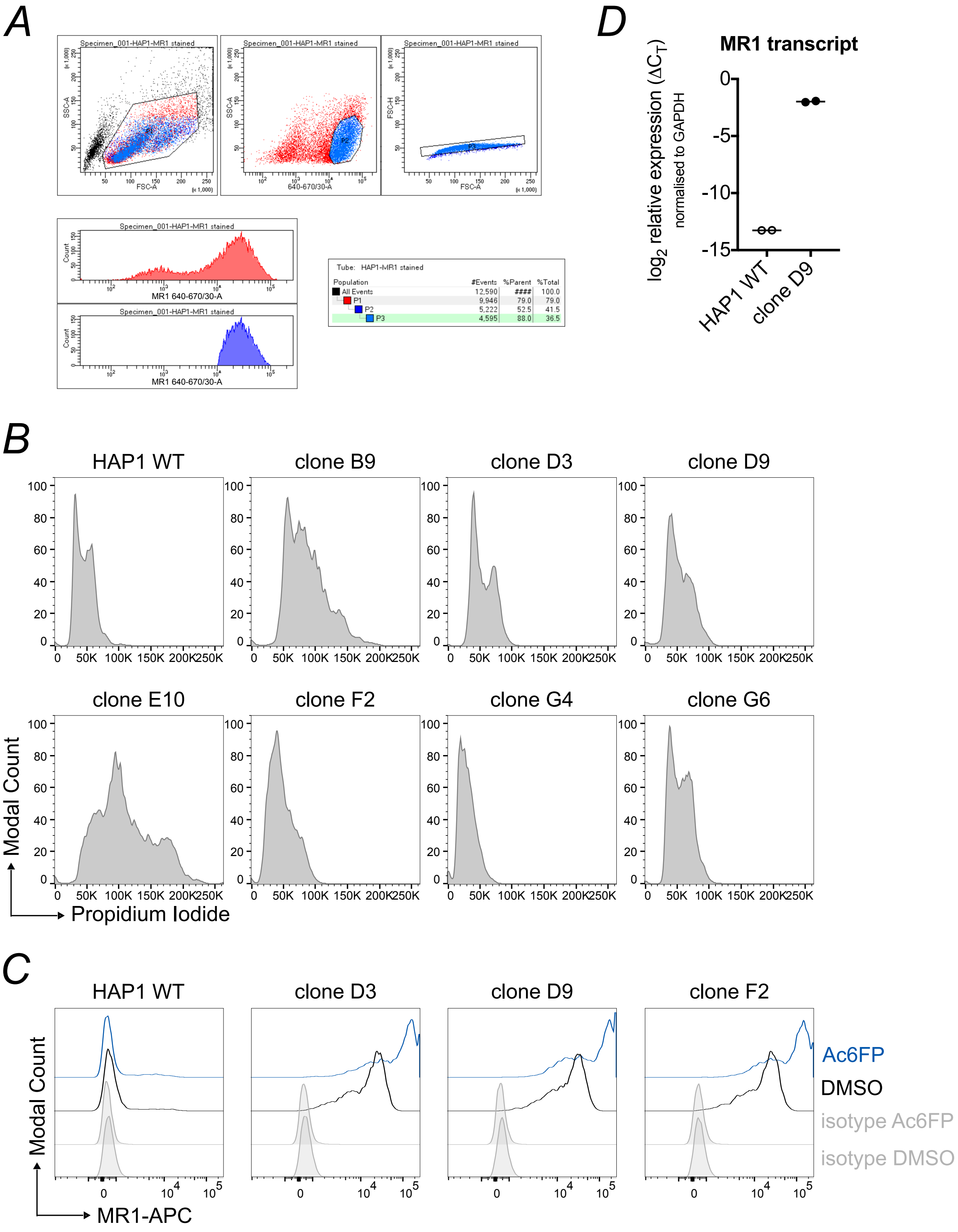

**Supporting Information Figure 1: HAP1 parent clones over-expressing MR1.** *A*, HAP1 WT cells were transduced with lentiviral particles encoding the human MR1 cDNA sequence, stained for surface expression of MR1, and single cell sorted to generate HAP1.MR1 parent clones. Sorted population: P3. *B* - *D*, clonal cell lines were screened for ploidy (*B*) and selected haploid clones were incubated with 10 µg/ml Ac6FP or the equivalent volume of solvent control DMSO for 5 h and stained for MR1 surface expression (*C*). Representative clones are shown in *B*. MR1 transcript levels were measured by qPCR in HAP1.MR1 parent clone D9 (*D*). Symbols in *D* represent technical replicates. Data in *C* + *D* are representative of two experiments.

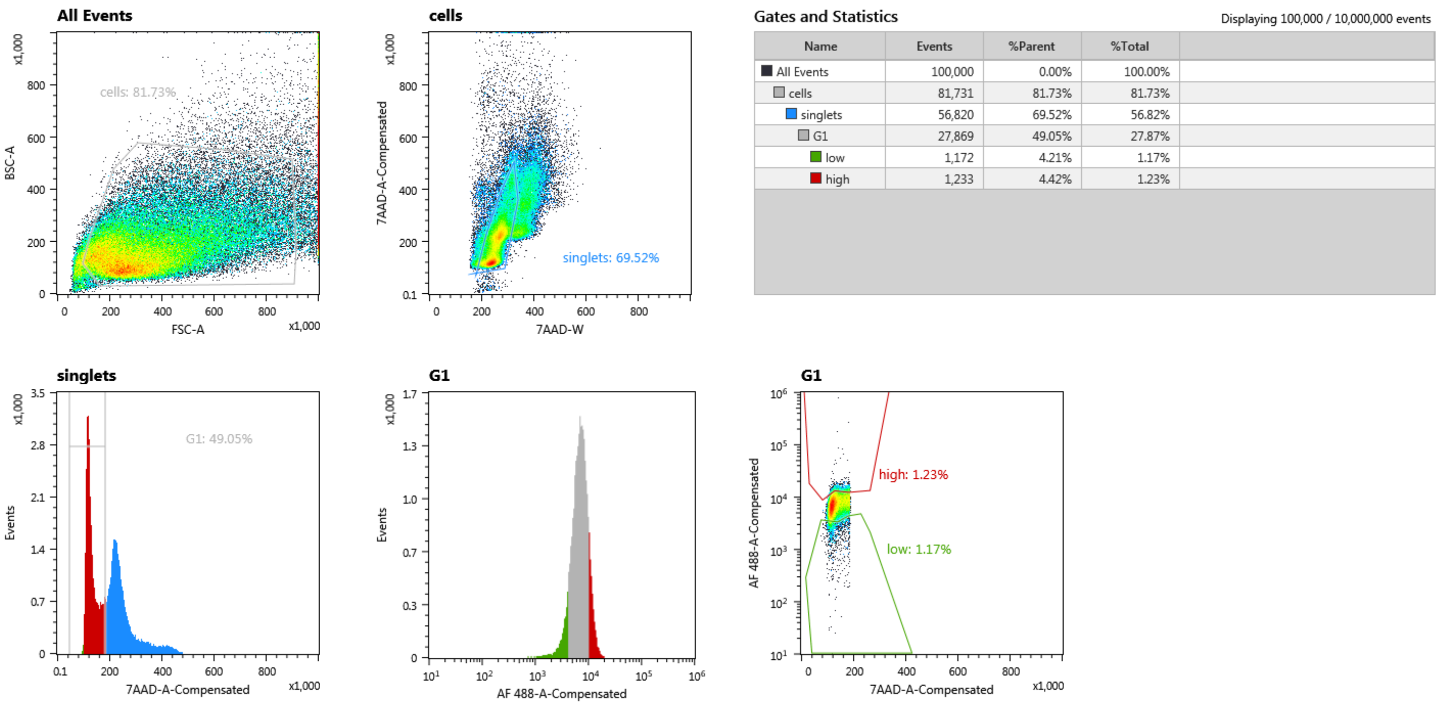

**Supporting Information Figure 2: Sorting strategy for gene-trap screen sort.** The mutagenized HAP1.MR1 clone D9 was stained as described in Materials and Methods and sorted as shown. Gates were set on the G1 population to include only cells haploid at the time of sorting.

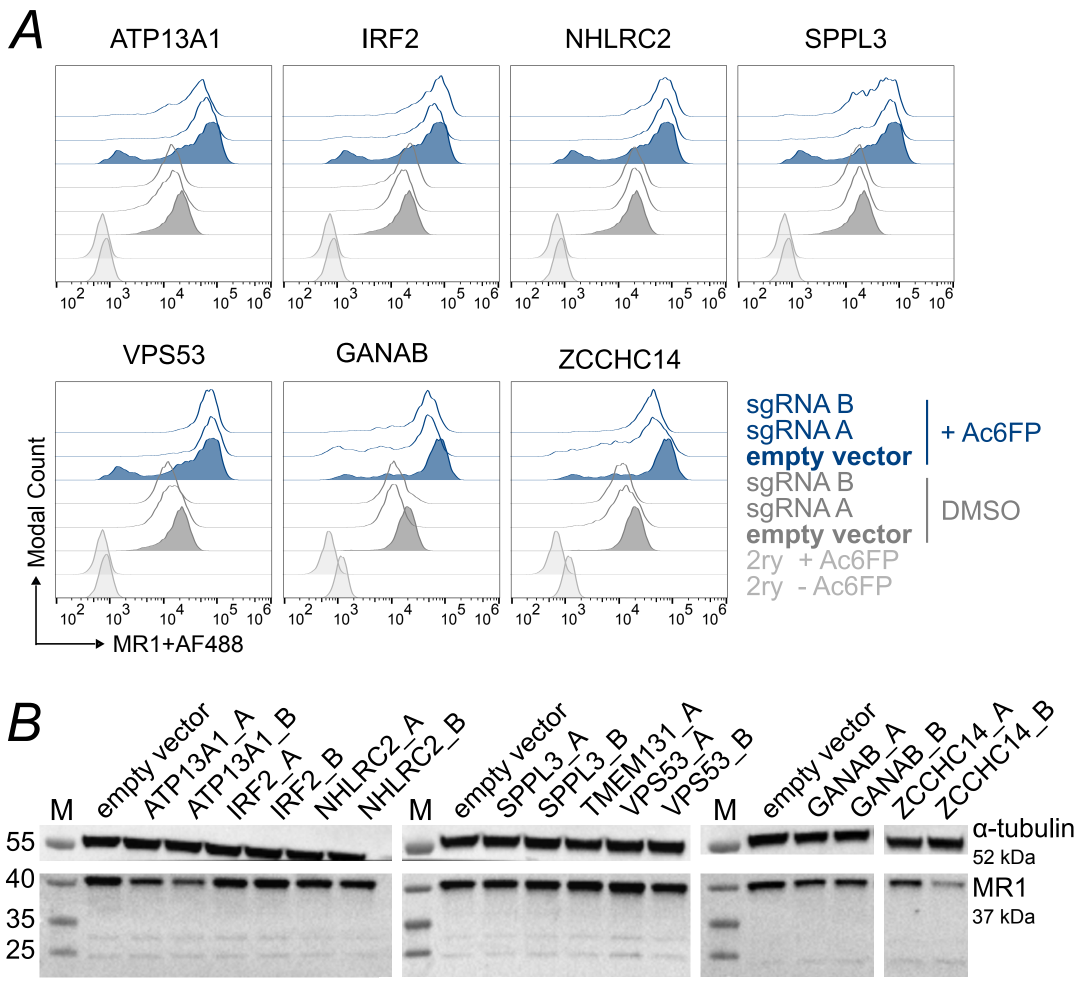

**Supporting Information Figure 3: Validation in bulk transfected HAP1.MR1 parent clone D9.** *A* + *B*, the HAP1.MR1 parent clone D9 was transfected with plasmids encoding the Cas9 protein and either no sgRNA (empty vector; filled histograms) or one of two different sgRNAs targeting the indicated gene (sgRNA A/B; empty histograms). Transfected cells were selected with puromycin and analyzed by flow cytometry (*A*) and Western Blot (*B*). *A*, cells were incubated with 5 µg/ml Ac6FP (blue) or DMSO (grey) for 5 h before surface staining with the anti-MR1 primary antibody (clone 26.5) and an AlexaFluor488-coupled secondary antibody (2ry). Histograms shown are gated on single cells in G1 phase based on propidium iodide staining. The same empty vector and secondary antibody controls are shown for cells transfected with sgRNAs targeting ATP13A1, IRF2, NHLRC, SPPL3, and VPS53 (batch 1) and GANAB and ZCCHC14 (batch2), respectively. *B*, The same number of cells was lysed and lysates were split and analyzed for MR1 (bottom) or loading control alpha-tubulin (top). Membranes were cut prior to primary antibody incubation. M in *B* denotes protein marker and molecular weights of visible marker bands are indicated in kDa to the left of the blots.

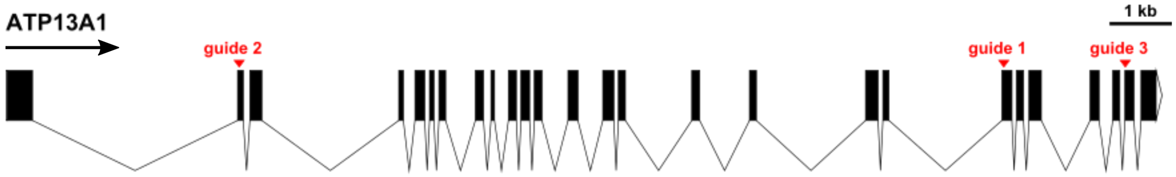

**Supporting Information Figure 4: Positions of the sgRNA target sequences with respect to the intron/exon structure of the ATP13A1 gene.** Arrow denotes direction of transcription. The plot was generated using the Exon-Intron Graphic Maker available at wormweb.org/exonintron based on the NCBI Reference Sequence NC_000019.10.

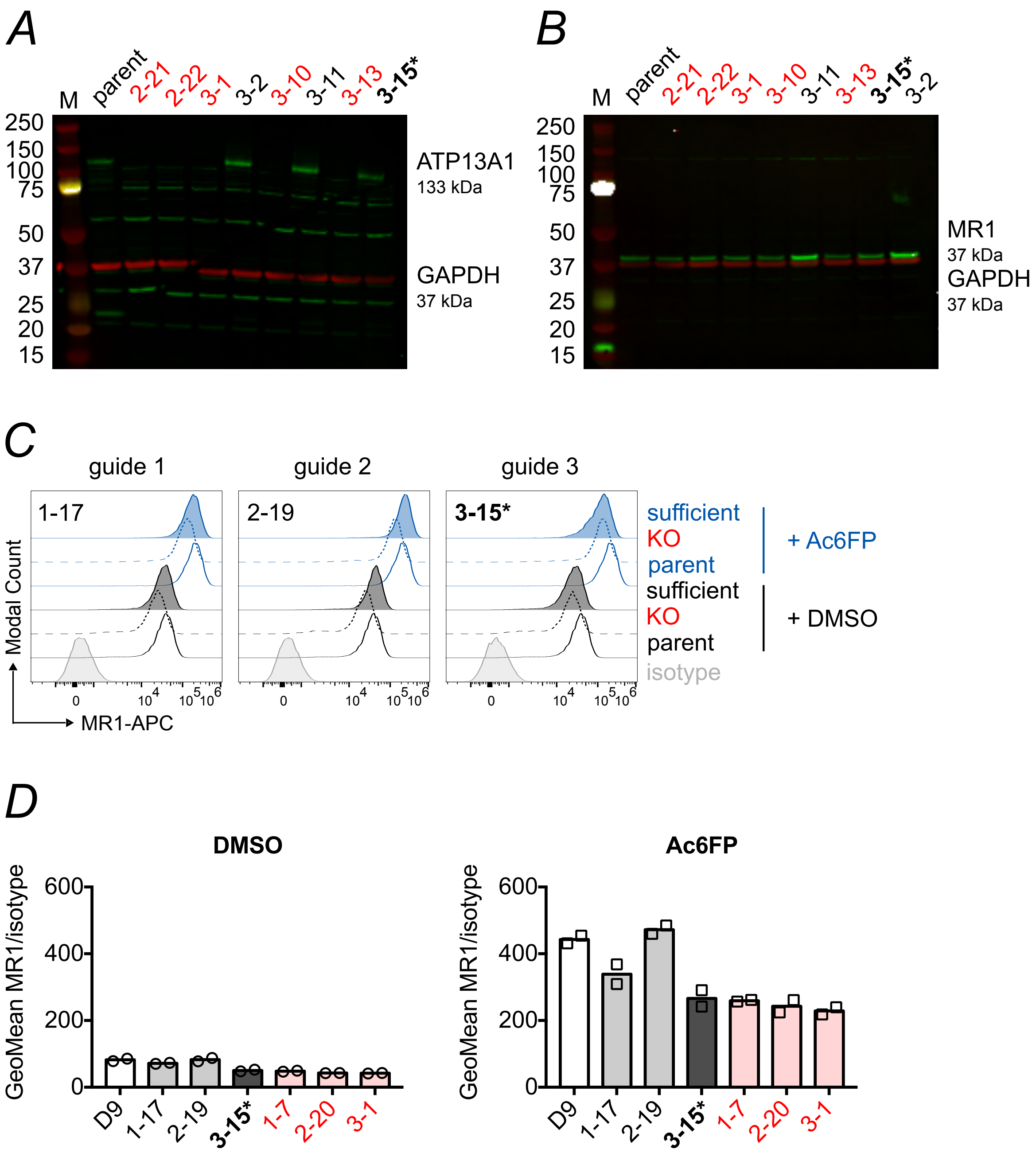

**Supporting Information Figure 5: HAP1.MR1 ATP13A1 CRISPR clone 3-15 expresses ATP13A1 but has a KO-like phenotype.** *A* + *B*, subset of HAP1.MR1 ATP13A1 CRISPR clones and the parent clone D9 (parent) were analysed for expression of ATP13A1 (*A*) and MR1 (*B*) by Western Blot. M denotes molecular weight marker and molecular weights of visible marker bands are indicated in kDa to the left of the blot. Western Blot of MR1 for clone 3-15 is representative of multiple experiments including the one shown in Supporting Information Figure 8. Lysates of the D9 parent clone were made prior to those of the clones. *C* + *D*, the D9 parent clone (empty histograms) and one KO (dashed lines) and one ATP13A1-expressing (filled histograms) clone per sgRNA were incubated with 5 µg/ml Ac6FP (blue) or the equivalent volume of DMSO (grey) for 5 h before MR1 surface expression was measured by flow cytometry using MR1 antibody clone 26.5. Representative histograms are shown in *C* and geometric mean of technical duplicates is shown in *D* with median. The same D9 parent clone samples are shown in each histogram for comparison. Both duplicates were normalised to the same isotype sample. Isotypes shown are those of the KO clones treated with Ac6FP. ATP13A1 KO clones are highlighted in red and the ATP13A1 mutant clone 3-15 is highlighted in bold and starred.

```
protein

no    ref   cttggccaatgcccctgagcgggttgtcgagcggcga-cggcggccccgggacagcccaa    1 bp insertion
                  ||||||||||||||||||||||||||||||||||||| ||||||||||||||||||||||    → frame shift
      1-7   CTTGGCCAATGCCCCTGAGCGGGTTGTCGAGCGGCGAGCGGCGGCCCCGGGACAGCCCAA

yes   ref   cttggccaatgcccctgagcgggttgtcgagcggcgacggcggccccgggacagcccaac    9 bp deletion
                  |||||||||||||||||||||||||||||||||||         ||||||||||||||||    → ERRRRPR to ERPR
      1-17  CTTGGCCAATGCCCCTGAGCGGGTTGTCGAGCGGC---------CCCGGGACAGCCCAAC    → 3 aa deleted in frame

yes   ref   cgaccccagcaaagcgaccttttgtgaaggtggtgccaacccccaacaatggctccacgga    2 bp swapped
                  |||||||||||||||||||||  |||||||||||||||||||||||||||||||||||||    → ATFVK to ATLLK
      2-19  CGACCCCAGCAAAGCGACCTTGTTGAAGGTGGTGCCAACCCCCAACAATGGCTCCACGGA    → 2 aa changed

no    ref   cgaccccagcaaagcgaccttttgtgaaggtggtgccaacccccaacaatggctccacgga    18 bp deletion
                  ||||||||||||||||                  ||||||||||||||||||||||||||    → ATFVKVVPT to APT
      2-20  CGACCCCAGCAAAGCG------------------CCAACCCCCAACAATGGCTCCACGGA    → 6 aa deleted in frame

no    ref   ccctcatccccacagggcccgcccttcatggagagcctgcccgagaacaagcccctggtg    2 bp deletion
                  ||||||||||||||||||||||  ||||||||||||||||||||||||||||||||||||    → frame shift
      3-1   CCCTCATCCCCACAGGGCCCGC--TTCATGGAGAGCCTGCCCGAGAACAAGCCCCTGGTG

yes   ref   ccctcatccccacagggcccgcccttcatggagagcctgcccgagaacaagcccctggtg    6 bp deletion
                  ||||||||||||||||||||||      ||||||||||||||||||||||||||||||||    → PFMESL to LESL
      3-15  CCCTCATCCCCACAGGGCCCGC------TGGAGAGCCTGCCCGAGAACAAGCCCCTGGTG    → conserved aa changed
```

**Supporting Information Figure 6: Sanger sequencing results of selected HAP1.MR1 ATP13A1 CRISPR clones.** Genomic regions around sgRNA target sites were amplified and sequenced in HAP1.MR1 ATP13A1 clones using the primers shown in Supporting Information Table 4. Names of the clones are given on the left and sequences are shown aligned to the NCBI Reference Sequence NC_000019.10 with sgRNA target sites highlighted in red. Consequences of gene editing are summarised to the right of each sequence. Protein expression based on Western Blot analysis is given to the left of each sequence. bp = base pair; aa = amino acid. Alignments generated with SerialCloner version 2.6.1.

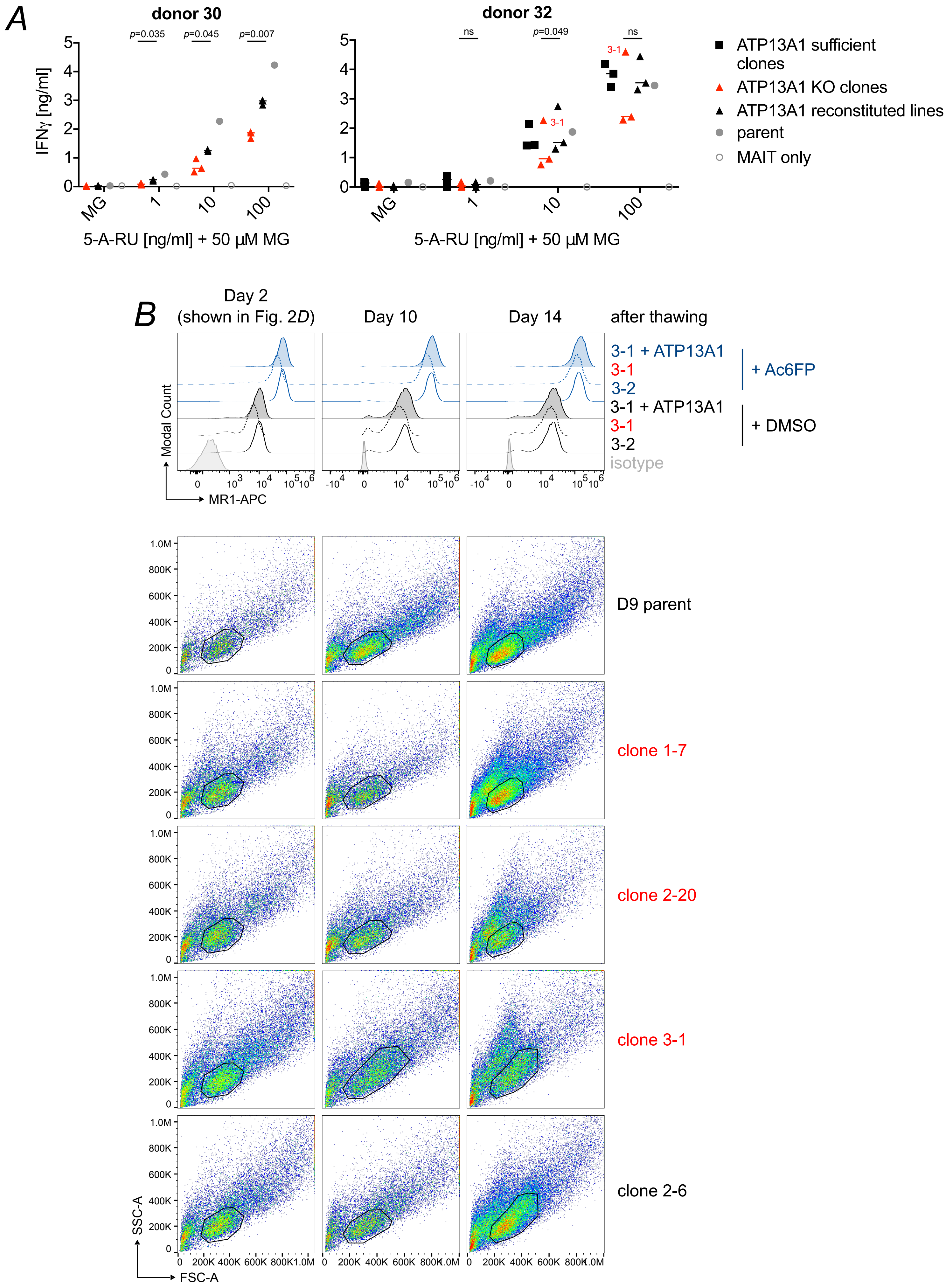

**Supporting Information Figure 7: Clone 3-1 changed morphology and lost the MR1 phenotype over time.** *A*, HAP1.MR1 ATP13A1 KO clones and their polyclonal reconstituted cell lines were pulsed with 5-A-RU+50 µM MG and incubated with sorted human MAIT cells for at least 36 h. IFNγ in the supernatants was quantified by ELISA. *(Legend continued on next page)*

*(Legend continued from previous page)* Each symbol represents the mean of technical duplicates for one clone or cell line and median is shown for the groups. Statistical significance of differences between ATP13A1 KO clones and reconstituted lines was analysed using two-tailed, paired *t* tests. The HAP1.MR1 parent clone D9 and ATP13A1 sufficient clones are shown for reference but were not included in statistical analyses. For details on ATP13A1 sufficient clones shown see Supporting Information Table 3. MAIT cell donors for each replicate experiment are shown above the plots. The experiment using MAIT cell donor 32 was performed with the same cells shown in *B* and carried out on day 3 after thawing. The experiment using donor 30 was performed several months earlier. *B,* three HAP1.MR1 ATP13A1 sufficient clones (solid lines), three HAP1.MR1 ATP13A1 KO clones (dashed lines), and their polyclonal reconstituted cell lines (filled; see Fig. 2*B-D*) were thawed and incubated with 5 µg/ml of Acetyl-6-formylpterin (Ac6FP, blue) or DMSO (black) for 5 h before staining for MR1 surface expression.The experiment was repeated three times and days after thawing are shown on top. Data for clones generated with sgRNA 3 is shown for all three experiments to illustrate that KO clone 3-1 lost the phenotype in the third experiment. Isotype control staining is from sufficient clone 3-2. Forward scatter (FSC) vs side scatter (SSC) plots for each experiment are also shown for selected clones to illustrate changes in morphology in KO clone 3-1 and sufficient clone 2-6. The same morphological change occurred in parallel in the polyclonal reconstituted cell line 3-1 + ATP13A1 (data not shown).  ATP13A1 KO clones are highlighted in red. ns = $p > 0.05$.

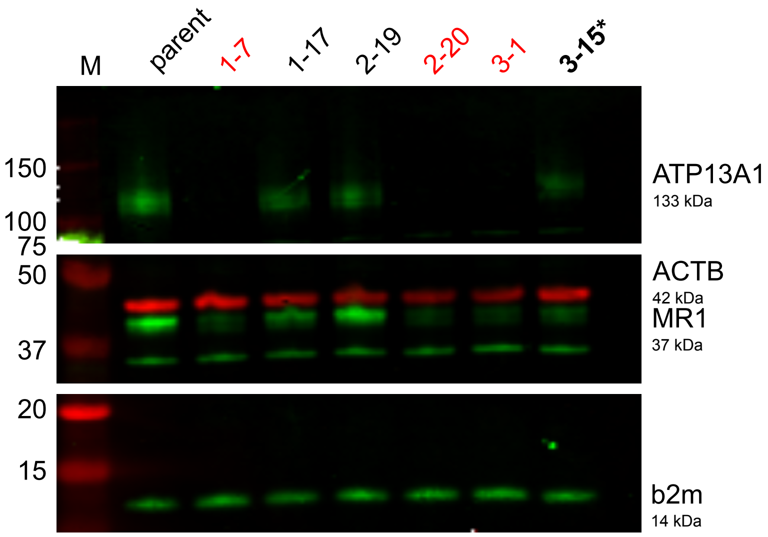

**Supporting Information Figure 8: Expression of β2-microglobulin is not a limiting factor in HAP1.MR1 ATP13A1 KO clones.** Protein expression of β2-microglobulin in HAP1.MR1 clones expressing ATP13A1 or not. Parent denotes HAP1.MR1 parent clone D9. Lysates were from the same experiment as those in Supporting Information Figure 14. M denotes protein marker and molecular weights of visible marker bands are indicated in kDa to the left of the blots. Loading control ACTB is shown in red and ATP13A1, MR1 and β2m are shown in green. The membrane was cut at the 25 kDa and 75 kDa marker bands prior to antibody incubation. Western Blot of β2m is representative of two independent experiments for KO clones 1-7, 2-20, and 3-1 but the ATP13A1 sufficient clones were not included in the other experiment. ATP13A1 KO clones are highlighted in red and the ATP13A1 mutant clone 3-15 is highlighted in bold and starred.

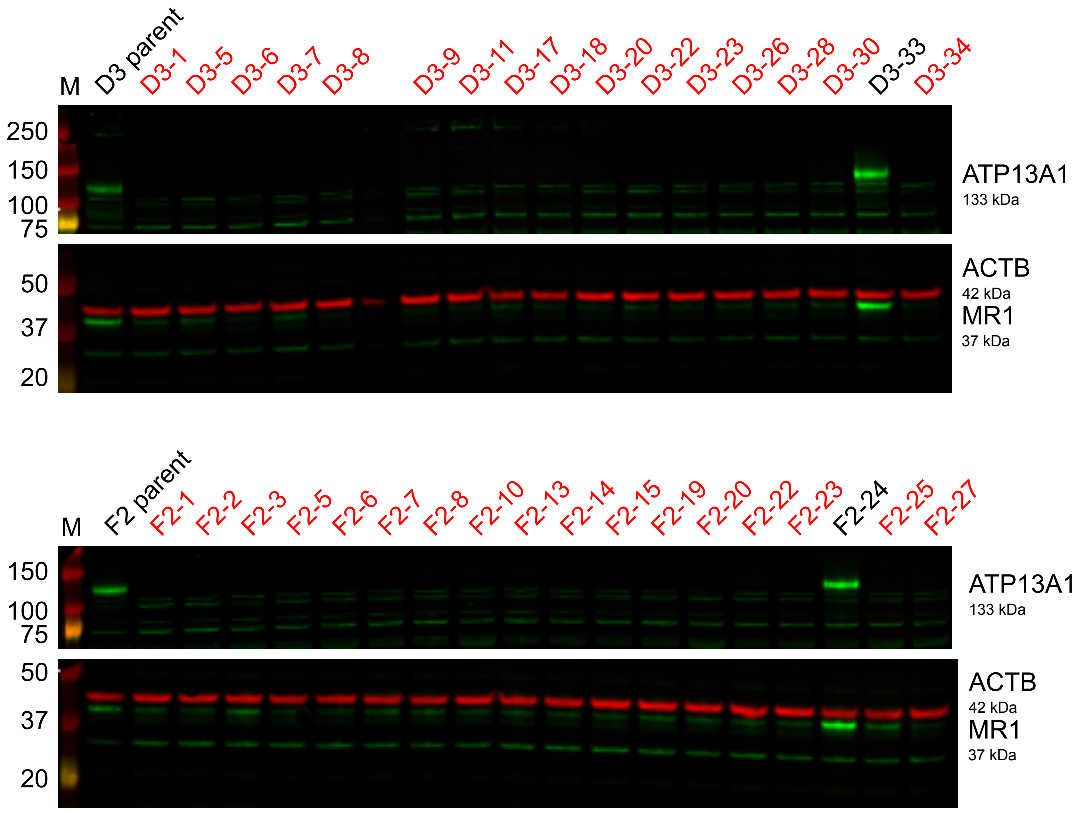

**Supporting Information Figure 9: Validation of total MR1 phenotype in HAP1.MR1 parent clones D3 and F2.** The HAP1.MR1 parent clones D3 and F2 (see Supporting Information Figure 1) were transfected with CRISPR/Cas9 plasmids encoding ATP13A1-targeting sgRNA2 and mRuby+ single cells were sorted. The generated clonal cell lines were analysed by Western Blot. Membranes were cut and probed with anti-ATP13A1 (top, green) or anti-MR1 (bottom, green) and anti-β actin (ACTB; bottom, red). Molecular weight of marker bands is shown in kDa to the left of the blots. Lysates of parent clones were prepared at a different time point than clones.

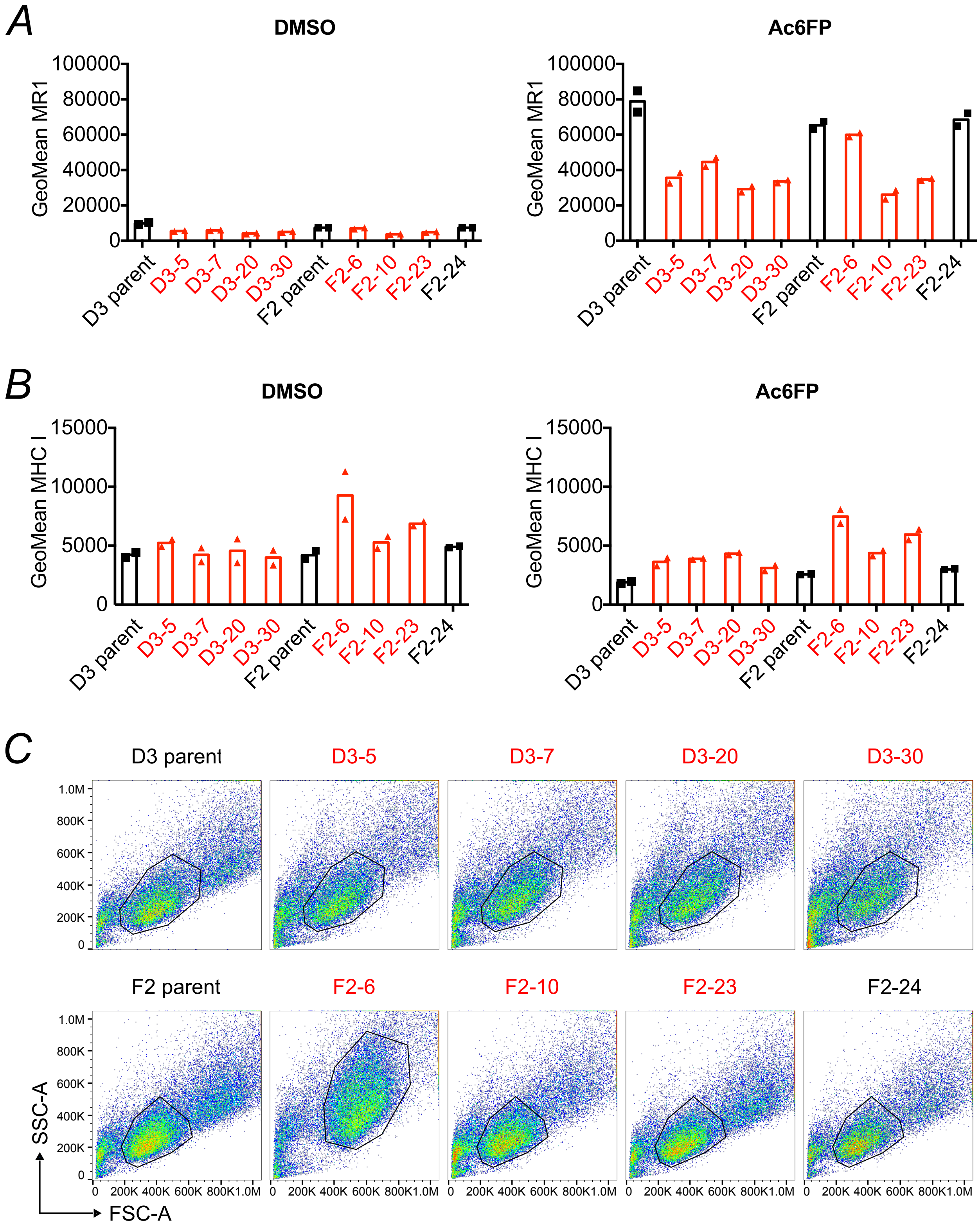

**Supporting Information Figure 10: Validation of surface MR1 phenotype in HAP1.MR1 parent clones D3 and F2.** The HAP1.MR1 parent clones D3 and F2 (see Supporting Information Figure 1) were transfected with CRISPR/Cas9 plasmids encoding ATP13A1-targeting sgRNA2 and mRuby+ single cells were sorted. *A*, a subset of the clones was incubated with 5 µg/ml Ac6FP (right) or the equivalent volume of DMSO (left) for 5 h before staining for MR1 (*A*) and MHC class I (*B*) surface expression for flow cytometry. Technical duplicates for each clone are shown. Forward scatter (FSC) vs side scatter (SSC) profiles are shown in (*C*). ATP13A1 KO clones are highlighted in red.

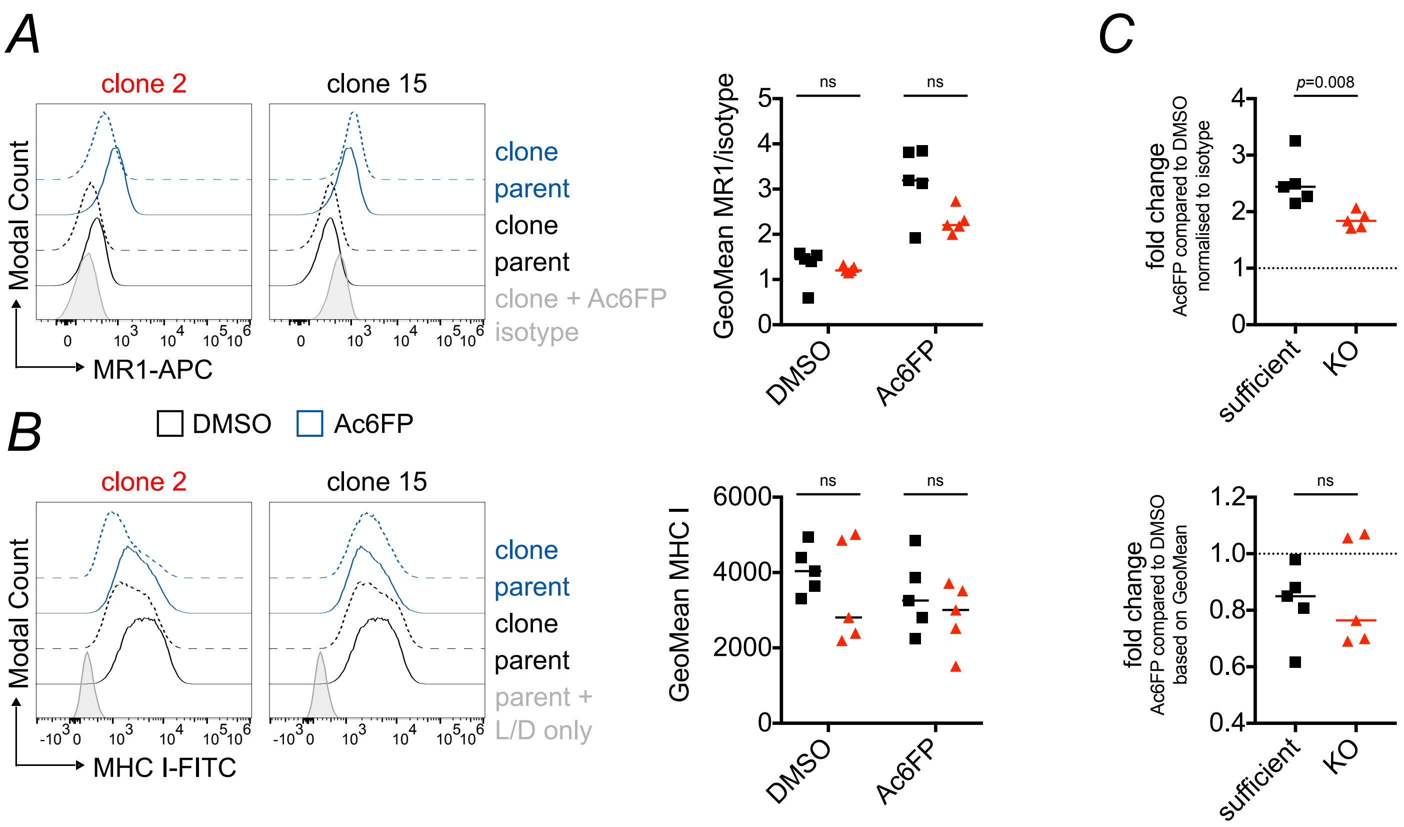

**Supporting Information Figure 11: Replicate of experiment shown in Figure 3*C* - *D* - MR1 surface expression is reduced in THP-1 ATP13A1 KO clones.** A THP-1 WT clone was transiently transfected with a CRISPR/Cas9 plasmid encoding sgRNA 2 targeting the first exon of ATP13A1 and cells were sorted and screened as shown in Figure 3. *A* + *B*, ten clones were incubated with Acetyl-6-formylpterin (Ac6FP, blue) or DMSO (black) for 5 h before staining for MR1 (*A*) and MHC class I (*B*) surface expression. Histograms for two representative clones are shown on the left and cumulative data from all 10 clones are shown on the right. The same parent sample is shown in both histograms for comparison. *C*, the same data as in *A* + *B* shown as fold-change of Geometric Mean Fluorescence Intensity (GeoMean) with Ac6FP compared to DMSO control. Results are supported by another experiment each although the repeat for *A* differed in statistical significance (see Figure 3*C* - *D*). Data are shown with median and statistical significance was calculated using the Mann-Whitney test. L/D = live/dead stain. ATP13A1 KO clones are highlighted in red. ns = $p > 0.05$.

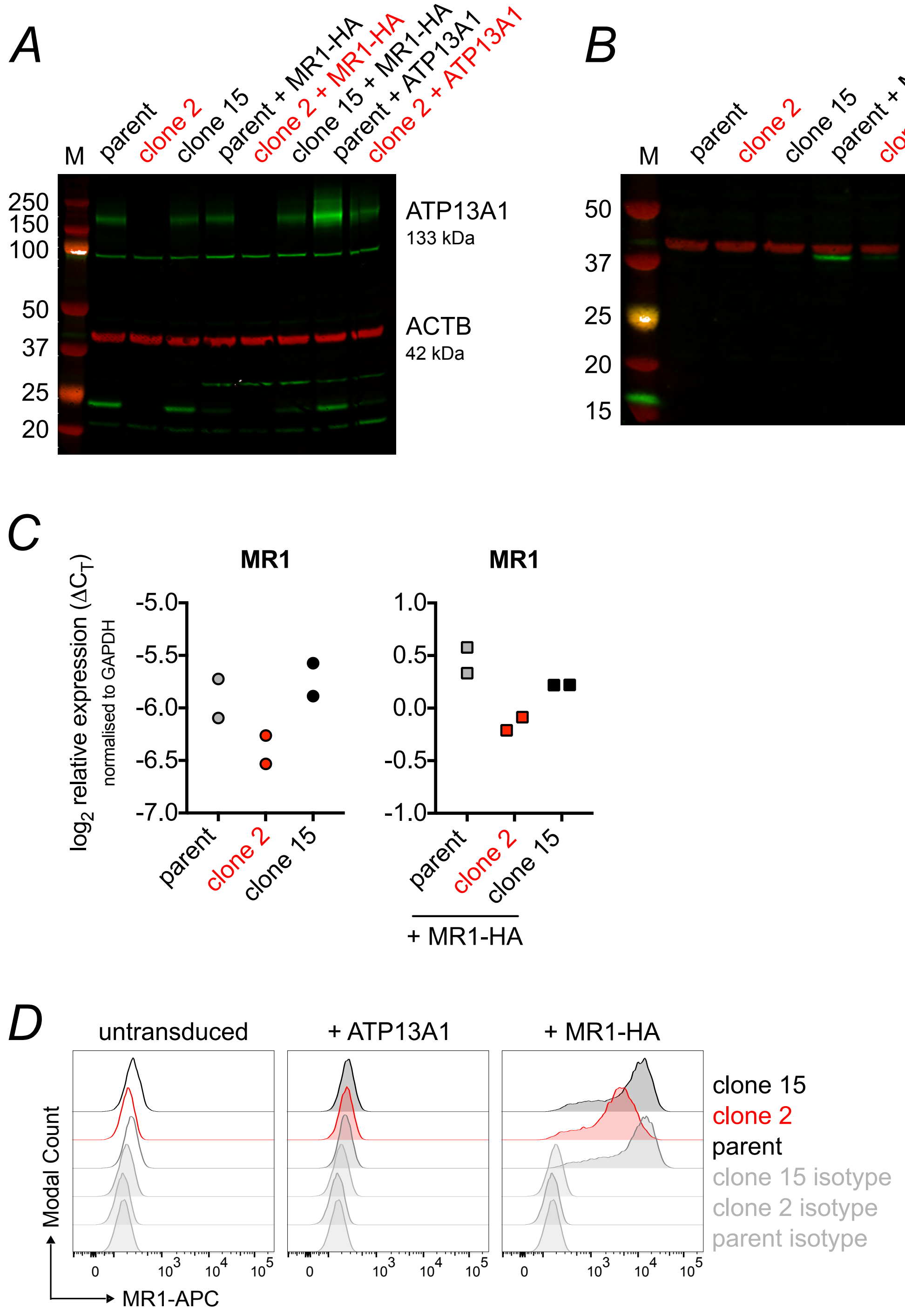

**Supporting Information Figure 12: Overexpression of ATP13A1 and MR1-HA in THP-1 clones.** The THP-1 parent clone, the THP-1 ATP13A1 KO clone 2, and the THP-1 ATP13A1 sufficient clone 15 were transduced with lentiviral particles encoding either ATP13A1 or an HA-tagged version of MR1 (MR1-HA). ATP13A1-overexpressing cells were selected with Blasticidin S. MR1-HA-expressing cells were sorted based on MR1 surface expression. The resulting cell lines were analysed by Western blot (*A* + *B*), qRT-PCR (*C*) and flow cytometry (*D*). Symbols in *C* represent technical duplicates. The same isotype controls are shown in all three plots in *D* as the samples were acquired together.

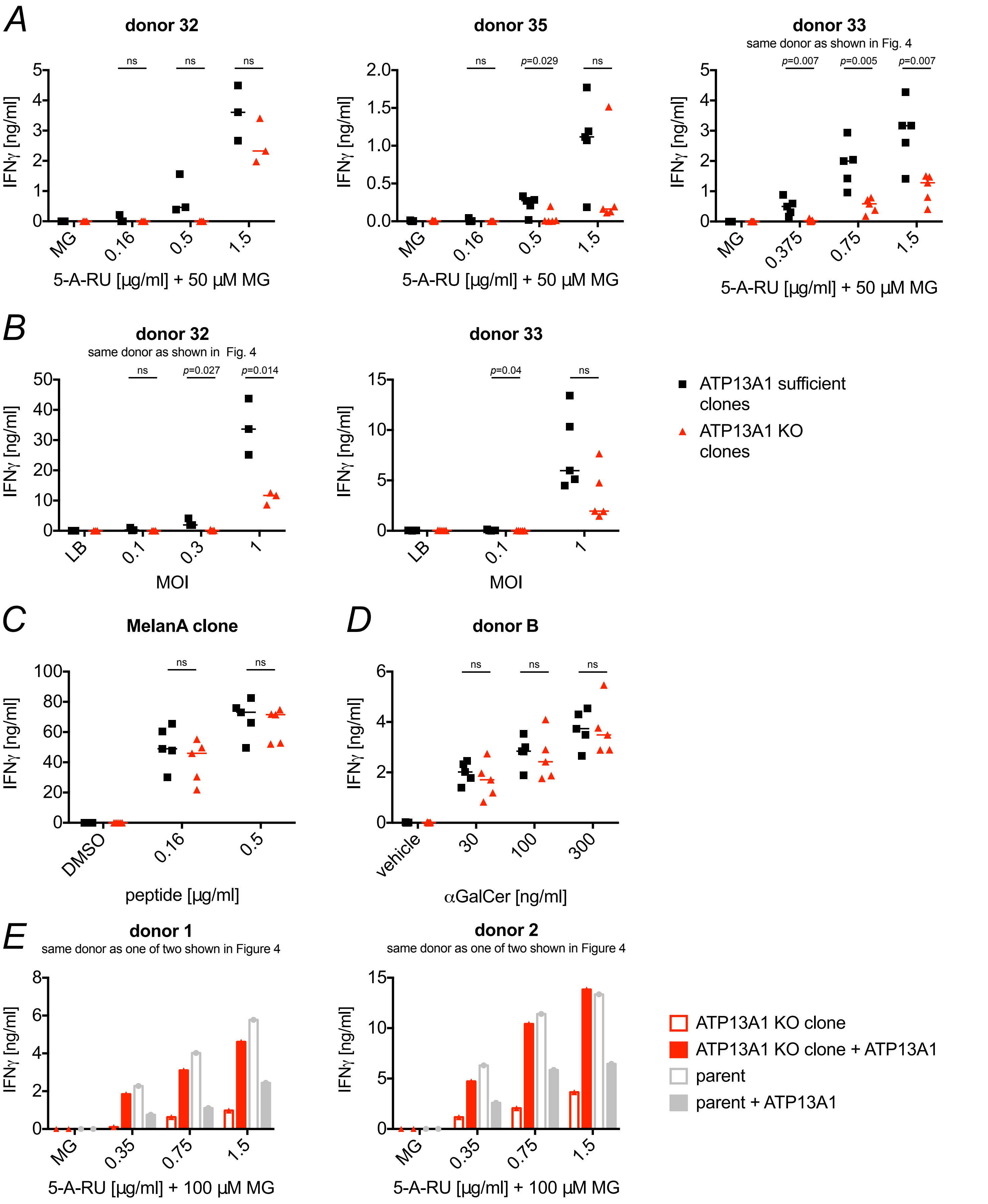

**Supporting Information Figure 13: Replicates of experiments shown in Figure 4 - MR1-mediated antigen presentation is reduced in THP-1 ATP13A1 KO clones.** *A – E*, THP-1 ATP13A1 sufficient or KO clones (*A-D*) or THP-1 parent and ATP13A1 KO clone 2 transduced to over-express ATP13A1 (*E*) were incubated with 5-A-RU and MG (*A* and *E*), *E.coli* (*B*), MelanA peptide (*C*), or αGalCer (*D*) before co-culture with sorted human MAIT cells (*A*, *B* + *E*), a MelanA-reactive T cell clone (*C*), or sorted human iNKT cells (*D*) as described in Figure 4. Each symbol represents the mean of technical duplicates for one clone and median is shown for the groups (*A – D*). ATP13A1 KO clones are highlighted in red. MAIT and iNKT cell donors for each experiment are indicated on top of the graphs where applicable. Statistical significance of differences between indicated groups was analysed using unpaired *t* tests. ns = $p > 0.05$.

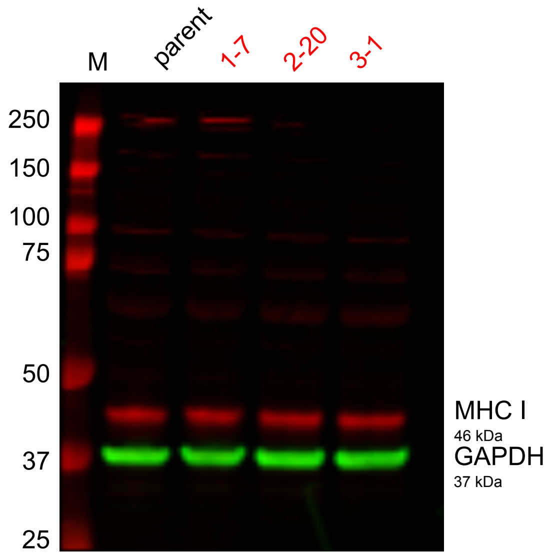

**Supporting Information Figure 14: MHC class I protein levels are unaffected in HAP1.MR1 ATP13A1 KO clones.** Cell lysates from DMSO-treated HAP1.MR1 parent clone D9 (parent) and the three independent ATP13A1 KO clones 1-7, 2-20, and 3-1 generated for the experiment shown in Figure 6A were separated by SDS-PAGE and immuno-blotted for MHC class I (red, HC10 antibody) and the loading control GAPDH (green). M denotes molecular weight marker and molecular weights of visible marker bands are indicated in kDa to the left of the blot. Western Blot is representative of two experiments.

**Supporting Information Table 1: Metadata of the screen.**

| Cells in MR1$^{hi}$ | Cells in MR1$^{low}$ | Unique Insertions in MR1$^{hi}$ | | Unique Insertions in MR1$^{low}$ | |
|---|---|---|---|---|---|
| | | sense | antisense | sense | antisense |
| 10.3x10$^{6}$ | 10.2x10$^{6}$ | 8.45x10$^{5}$ | 9.27x10$^{5}$ | 8.98x10$^{5}$ | 9.68x10$^{5}$ |

**Supporting Information Table 2: Gene-trap screen hits.** The number of insertions in sense orientation in the $MR1^{hi}$ fraction (high.sense), the number of insertions in the $MR1^{low}$ fraction (low.sense), the false discovery rate-corrected p-value (fcpv), and the mutation index (MI, see Materials and Methods) are shown for each statistically significant (fcpv < 0.1) enriched genes identified in the gene-trap screen. ins = insertions. Sorted by fcpv.

| | high.sense | low.sense | padjust.sense | MI.sense | log2MI.sense | log10 ins |
|---|---|---|---|---|---|---|
| SPEN | 60 | 1051 | 4.41E-218 | 0.061 | -4.045 | 3.05E+00 |
| ZCCHC14 | 11 | 294 | 3.10E-65 | 0.040 | -4.653 | 2.48E+00 |
| DYRK1A | 115 | 561 | 5.74E-62 | 0.218 | -2.200 | 2.83E+00 |
| SETD1B | 27 | 324 | 1.49E-58 | 0.089 | -3.498 | 2.55E+00 |
| CUL3 | 368 | 90 | 2.94E-41 | 4.345 | 2.119 | 2.66E+00 |
| B2M | 5 | 155 | 1.47E-34 | 0.034 | -4.867 | 2.20E+00 |
| PPP2R4 | 320 | 86 | 1.12E-32 | 3.954 | 1.983 | 2.61E+00 |
| ZNF292 | 222 | 568 | 3.84E-28 | 0.415 | -1.269 | 2.90E+00 |
| TIPRL | 159 | 18 | 4.28E-28 | 9.386 | 3.230 | 2.25E+00 |
| PARN | 115 | 379 | 1.83E-27 | 0.322 | -1.634 | 2.69E+00 |
| KMT2A | 5 | 128 | 2.23E-27 | 0.041 | -4.591 | 2.12E+00 |
| TMEM131 | 324 | 111 | 4.77E-25 | 3.102 | 1.633 | 2.64E+00 |
| RALGAPB | 208 | 48 | 4.05E-24 | 4.604 | 2.203 | 2.41E+00 |
| ACACA | 380 | 774 | 1.31E-23 | 0.521 | -0.940 | 3.06E+00 |
| GANAB | 7 | 118 | 7.46E-23 | 0.063 | -3.988 | 2.10E+00 |
| DOT1L | 26 | 169 | 3.19E-22 | 0.163 | -2.613 | 2.29E+00 |
| HDAC9 | 4323 | 3662 | 4.29E-21 | 1.255 | 0.328 | 3.90E+00 |
| PPP4R2 | 173 | 40 | 6.67E-20 | 4.595 | 2.200 | 2.33E+00 |
| WAC | 39 | 181 | 1.82E-18 | 0.229 | -2.127 | 2.34E+00 |
| VPS29 | 24 | 145 | 7.04E-18 | 0.176 | -2.508 | 2.23E+00 |
| MLLT10 | 218 | 470 | 4.44E-16 | 0.493 | -1.021 | 2.84E+00 |
| SND1 | 40 | 165 | 7.38E-15 | 0.258 | -1.957 | 2.31E+00 |
| MR1 | 28 | 137 | 2.90E-14 | 0.217 | -2.204 | 2.22E+00 |
| REPS1 | 47 | 175 | 4.70E-14 | 0.285 | -1.810 | 2.35E+00 |
| HTT | 63 | 198 | 6.28E-13 | 0.338 | -1.565 | 2.42E+00 |
| ZCCHC11 | 285 | 135 | 7.65E-13 | 2.243 | 1.166 | 2.62E+00 |
| CREBBP | 93 | 246 | 1.87E-12 | 0.402 | -1.316 | 2.53E+00 |
| EP300 | 40 | 152 | 2.08E-12 | 0.280 | -1.839 | 2.28E+00 |
| LIN28B | 338 | 179 | 6.35E-12 | 2.006 | 1.005 | 2.71E+00 |
| ATF7IP | 505 | 310 | 6.40E-12 | 1.731 | 0.792 | 2.91E+00 |
| ANKRD17 | 329 | 173 | 9.42E-12 | 2.021 | 1.015 | 2.70E+00 |
| DBF4 | 102 | 25 | 9.59E-11 | 4.335 | 2.116 | 2.10E+00 |
| PICALM | 294 | 153 | 1.29E-10 | 2.042 | 1.030 | 2.65E+00 |
| ZBTB20 | 4972 | 4550 | 1.47E-10 | 1.162 | 0.216 | 3.98E+00 |
| IRF2 | 14 | 86 | 2.39E-10 | 0.173 | -2.532 | 2.00E+00 |
| NCOA2 | 736 | 523 | 6.84E-10 | 1.495 | 0.581 | 3.10E+00 |
| PPP6R3 | 247 | 123 | 8.04E-10 | 2.134 | 1.093 | 2.57E+00 |
| ERC1 | 359 | 605 | 9.43E-10 | 0.630 | -0.666 | 2.98E+00 |
| CARM1 | 158 | 325 | 1.25E-09 | 0.516 | -0.953 | 2.68E+00 |
| ROBO1 | 4089 | 3712 | 1.48E-09 | 1.171 | 0.228 | 3.89E+00 |
| ATP13A1 | 3 | 52 | 2.28E-09 | 0.061 | -4.028 | 1.74E+00 |
| ADSS | 19 | 92 | 2.44E-09 | 0.219 | -2.188 | 2.05E+00 |
| RQCD1 | 88 | 21 | 2.50E-09 | 4.452 | 2.155 | 2.04E+00 |
| TFDP1 | 81 | 18 | 3.82E-09 | 4.781 | 2.257 | 2.00E+00 |

| | high.sense | low.sense | padjust.sense | MI.sense | log2MI.sense | log10 ins |
|---|---|---|---|---|---|---|
| CDC25A | 214 | 104 | 5.39E-09 | 2.186 | 1.129 | 2.50E+00 |
| MNT | 117 | 257 | 5.85E-09 | 0.484 | -1.048 | 2.57E+00 |
| MGA | 127 | 272 | 6.27E-09 | 0.496 | -1.012 | 2.60E+00 |
| UBE2H | 571 | 865 | 1.23E-08 | 0.701 | -0.512 | 3.16E+00 |
| YWHAE | 50 | 147 | 1.34E-08 | 0.361 | -1.469 | 2.29E+00 |
| EIF4B | 102 | 231 | 1.43E-08 | 0.469 | -1.092 | 2.52E+00 |
| BAZ1B | 302 | 176 | 3.66E-08 | 1.823 | 0.866 | 2.68E+00 |
| KDM3B | 119 | 253 | 3.93E-08 | 0.500 | -1.001 | 2.57E+00 |
| ACTR3 | 52 | 147 | 4.31E-08 | 0.376 | -1.412 | 2.30E+00 |
| USP14 | 755 | 1084 | 5.24E-08 | 0.740 | -0.435 | 3.26E+00 |
| GPBP1 | 58 | 157 | 5.35E-08 | 0.392 | -1.350 | 2.33E+00 |
| DPH5 | 17 | 80 | 8.39E-08 | 0.226 | -2.147 | 1.99E+00 |
| CSNK2A1 | 20 | 84 | 2.14E-07 | 0.253 | -1.983 | 2.02E+00 |
| DACH1 | 2695 | 2412 | 2.48E-07 | 1.188 | 0.248 | 3.71E+00 |
| NAA30 | 63 | 13 | 2.73E-07 | 5.149 | 2.364 | 1.88E+00 |
| BIRC6 | 194 | 351 | 4.38E-07 | 0.587 | -0.768 | 2.74E+00 |
| SSR3 | 6 | 51 | 4.41E-07 | 0.125 | -3.000 | 1.76E+00 |
| FANCL | 65 | 15 | 4.74E-07 | 4.604 | 2.203 | 1.90E+00 |
| AHCY | 27 | 96 | 4.88E-07 | 0.299 | -1.743 | 2.09E+00 |
| XPO6 | 221 | 387 | 5.23E-07 | 0.607 | -0.721 | 2.78E+00 |
| RFXAP | 11 | 62 | 6.55E-07 | 0.188 | -2.408 | 1.86E+00 |
| DCAF7 | 6 | 50 | 7.20E-07 | 0.127 | -2.972 | 1.75E+00 |
| CDK6 | 344 | 220 | 7.47E-07 | 1.661 | 0.732 | 2.75E+00 |
| KMT2E | 416 | 281 | 8.83E-07 | 1.573 | 0.654 | 2.84E+00 |
| PUM1 | 54 | 140 | 1.35E-06 | 0.410 | -1.287 | 2.29E+00 |
| ARPC2 | 17 | 73 | 1.90E-06 | 0.247 | -2.015 | 1.95E+00 |
| RAB10 | 33 | 103 | 1.90E-06 | 0.340 | -1.555 | 2.13E+00 |
| VHL | 3 | 39 | 2.09E-06 | 0.082 | -3.613 | 1.62E+00 |
| TLK2 | 143 | 272 | 2.20E-06 | 0.558 | -0.841 | 2.62E+00 |
| ARHGEF38 | 194 | 105 | 3.32E-06 | 1.963 | 0.973 | 2.48E+00 |
| BPTF | 27 | 91 | 3.46E-06 | 0.315 | -1.666 | 2.07E+00 |
| RFWD2 | 212 | 365 | 3.48E-06 | 0.617 | -0.697 | 2.76E+00 |
| DCPS | 10 | 56 | 3.82E-06 | 0.190 | -2.398 | 1.82E+00 |
| IMMT | 10 | 56 | 3.82E-06 | 0.190 | -2.398 | 1.82E+00 |
| KIAA1033 | 102 | 210 | 4.52E-06 | 0.516 | -0.955 | 2.49E+00 |
| CUX1 | 122 | 238 | 5.11E-06 | 0.545 | -0.877 | 2.56E+00 |
| ASCC3 | 38 | 109 | 6.06E-06 | 0.370 | -1.433 | 2.17E+00 |
| ADNP | 166 | 86 | 6.27E-06 | 2.051 | 1.036 | 2.40E+00 |
| VAC14 | 60 | 15 | 7.39E-06 | 4.250 | 2.087 | 1.88E+00 |
| TAPBP | 99 | 39 | 9.12E-06 | 2.697 | 1.431 | 2.14E+00 |
| PMVK | 16 | 67 | 1.07E-05 | 0.254 | -1.979 | 1.92E+00 |
| NHLRC2 | 15 | 65 | 1.10E-05 | 0.245 | -2.028 | 1.90E+00 |
| HSD17B12 | 25 | 84 | 1.33E-05 | 0.316 | -1.661 | 2.04E+00 |
| NFYC | 95 | 38 | 2.20E-05 | 2.656 | 1.409 | 2.12E+00 |
| KDM2A | 156 | 278 | 2.68E-05 | 0.596 | -0.746 | 2.64E+00 |
| MOGS | 3 | 34 | 3.23E-05 | 0.094 | -3.415 | 1.57E+00 |
| TIMM50 | 3 | 34 | 3.23E-05 | 0.094 | -3.415 | 1.57E+00 |
| ALG8 | 67 | 150 | 3.29E-05 | 0.474 | -1.076 | 2.34E+00 |

| | high.sense | low.sense | padjust.sense | MI.sense | log2MI.sense | log10 ins |
|---|---|---|---|---|---|---|
| FBXO42 | 26 | 83 | 3.81E-05 | 0.333 | -1.587 | 2.04E+00 |
| RBM15 | 12 | 55 | 4.31E-05 | 0.232 | -2.109 | 1.83E+00 |
| TSC1 | 102 | 45 | 5.97E-05 | 2.408 | 1.268 | 2.17E+00 |
| ZNF236 | 31 | 90 | 6.17E-05 | 0.366 | -1.450 | 2.08E+00 |
| SRD5A3 | 57 | 132 | 6.55E-05 | 0.459 | -1.124 | 2.28E+00 |
| PBRM1 | 648 | 510 | 6.74E-05 | 1.350 | 0.433 | 3.06E+00 |
| CCDC6 | 32 | 91 | 7.62E-05 | 0.374 | -1.421 | 2.09E+00 |
| RAD51B | 198 | 118 | 7.69E-05 | 1.783 | 0.834 | 2.50E+00 |
| PPAT | 43 | 109 | 8.57E-05 | 0.419 | -1.255 | 2.18E+00 |
| VPS41 | 348 | 244 | 9.56E-05 | 1.515 | 0.600 | 2.77E+00 |
| NLGN1 | 1722 | 1536 | 9.84E-05 | 1.191 | 0.253 | 3.51E+00 |
| KDM6A | 143 | 76 | 0.000101732 | 1.999 | 0.999 | 2.34E+00 |
| SLTM | 107 | 204 | 0.000105749 | 0.557 | -0.844 | 2.49E+00 |
| RPTOR | 44 | 109 | 0.000136931 | 0.429 | -1.222 | 2.18E+00 |
| NDUFAF6 | 135 | 71 | 0.000136931 | 2.020 | 1.014 | 2.31E+00 |
| LAMTOR2 | 34 | 5 | 0.000138817 | 7.224 | 2.853 | 1.59E+00 |
| MTF2 | 71 | 150 | 0.000154636 | 0.503 | -0.992 | 2.34E+00 |
| YTHDF2 | 26 | 78 | 0.00015945 | 0.354 | -1.498 | 2.02E+00 |
| LAMTOR5 | 71 | 26 | 0.000164375 | 2.901 | 1.537 | 1.99E+00 |
| TTI1 | 2 | 28 | 0.000166392 | 0.076 | -3.720 | 1.48E+00 |
| ZFX | 144 | 252 | 0.000182205 | 0.607 | -0.720 | 2.60E+00 |
| NPM1 | 4 | 33 | 0.000217127 | 0.129 | -2.957 | 1.57E+00 |
| UBE2G2 | 15 | 56 | 0.000328165 | 0.285 | -1.813 | 1.85E+00 |
| CNOT2 | 54 | 122 | 0.000328165 | 0.470 | -1.089 | 2.25E+00 |
| GART | 74 | 151 | 0.000356901 | 0.521 | -0.942 | 2.35E+00 |
| TAP1 | 59 | 20 | 0.000356901 | 3.134 | 1.648 | 1.90E+00 |
| NFIC | 121 | 218 | 0.000361603 | 0.590 | -0.762 | 2.53E+00 |
| CTNNA1 | 347 | 510 | 0.000370509 | 0.723 | -0.469 | 2.93E+00 |
| TRPM3 | 1769 | 1599 | 0.000390892 | 1.176 | 0.233 | 3.53E+00 |
| VPS53 | 5 | 34 | 0.000454842 | 0.156 | -2.678 | 1.59E+00 |
| BRIP1 | 179 | 109 | 0.000500343 | 1.745 | 0.803 | 2.46E+00 |
| EIF3L | 9 | 43 | 0.000591275 | 0.222 | -2.169 | 1.72E+00 |
| RHEB | 4 | 31 | 0.000607614 | 0.137 | -2.867 | 1.54E+00 |
| UBE2L3 | 129 | 70 | 0.000611315 | 1.958 | 0.969 | 2.30E+00 |
| KAT7 | 16 | 57 | 0.000671761 | 0.298 | -1.746 | 1.86E+00 |
| LAMTOR4 | 65 | 25 | 0.000842366 | 2.762 | 1.466 | 1.95E+00 |
| SLIT2 | 1760 | 1601 | 0.000891219 | 1.168 | 0.224 | 3.53E+00 |
| SEC63 | 9 | 42 | 0.000923509 | 0.228 | -2.135 | 1.71E+00 |
| BMPR2 | 79 | 154 | 0.000930195 | 0.545 | -0.876 | 2.37E+00 |
| FIG4 | 45 | 13 | 0.001047423 | 3.678 | 1.879 | 1.76E+00 |
| RANBP1 | 48 | 108 | 0.001073813 | 0.472 | -1.083 | 2.19E+00 |
| MAGI2 | 1977 | 1818 | 0.001098718 | 1.156 | 0.209 | 3.58E+00 |
| C16orf62 | 36 | 89 | 0.001120299 | 0.430 | -1.219 | 2.10E+00 |
| POU6F2 | 1707 | 1554 | 0.001283363 | 1.167 | 0.223 | 3.51E+00 |
| PDS5A | 26 | 72 | 0.00128394 | 0.384 | -1.382 | 1.99E+00 |
| SGF29 | 6 | 35 | 0.001323309 | 0.182 | -2.457 | 1.61E+00 |
| TAF11 | 6 | 35 | 0.001323309 | 0.182 | -2.457 | 1.61E+00 |
| NUP188 | 10 | 43 | 0.001367396 | 0.247 | -2.017 | 1.72E+00 |

| | high.sense | low.sense | padjust.sense | MI.sense | log2MI.sense | log10 ins |
|---|---|---|---|---|---|---|
| ADK | 954 | 1224 | 0.001386462 | 0.828 | -0.273 | 3.34E+00 |
| GPC3 | 1634 | 1484 | 0.001386462 | 1.170 | 0.227 | 3.49E+00 |
| GTF3C2 | 16 | 54 | 0.001393145 | 0.315 | -1.668 | 1.85E+00 |
| DPH6 | 285 | 423 | 0.001393145 | 0.716 | -0.483 | 2.85E+00 |
| TAP2 | 46 | 14 | 0.001393145 | 3.491 | 1.804 | 1.78E+00 |
| NONO | 27 | 73 | 0.001500965 | 0.393 | -1.348 | 2.00E+00 |
| DLG2 | 4047 | 3900 | 0.001500965 | 1.103 | 0.141 | 3.90E+00 |
| UBE2G1 | 289 | 427 | 0.00153179 | 0.719 | -0.476 | 2.85E+00 |
| GNG7 | 213 | 330 | 0.001832058 | 0.686 | -0.544 | 2.73E+00 |
| TFAP4 | 83 | 157 | 0.001963272 | 0.562 | -0.832 | 2.38E+00 |
| FCHO2 | 106 | 56 | 0.002003223 | 2.011 | 1.008 | 2.21E+00 |
| EIF2A | 88 | 163 | 0.002262804 | 0.574 | -0.802 | 2.40E+00 |
| DNAJC7 | 55 | 115 | 0.00234994 | 0.508 | -0.977 | 2.23E+00 |
| UBR5 | 26 | 70 | 0.002401317 | 0.395 | -1.342 | 1.98E+00 |
| ZFR | 23 | 65 | 0.002558806 | 0.376 | -1.412 | 1.94E+00 |
| MTHFD1 | 25 | 68 | 0.002753731 | 0.391 | -1.356 | 1.97E+00 |
| FLCN | 37 | 10 | 0.002912256 | 3.931 | 1.975 | 1.67E+00 |
| WHSC1 | 102 | 181 | 0.002988261 | 0.599 | -0.740 | 2.45E+00 |
| VPS50 | 37 | 87 | 0.003088562 | 0.452 | -1.146 | 2.09E+00 |
| GGNBP2 | 96 | 172 | 0.003114704 | 0.593 | -0.754 | 2.43E+00 |
| VPS13D | 23 | 64 | 0.00360208 | 0.382 | -1.389 | 1.94E+00 |
| PSIP1 | 17 | 53 | 0.003840712 | 0.341 | -1.553 | 1.85E+00 |
| EXOC4 | 470 | 641 | 0.003840712 | 0.779 | -0.361 | 3.05E+00 |
| EZH2 | 35 | 83 | 0.004203 | 0.448 | -1.159 | 2.07E+00 |
| STT3B | 64 | 126 | 0.004295498 | 0.540 | -0.890 | 2.28E+00 |
| TASP1 | 234 | 164 | 0.004523555 | 1.516 | 0.600 | 2.60E+00 |
| CSMD3 | 1204 | 1078 | 0.004651848 | 1.187 | 0.247 | 3.36E+00 |
| BMP2K | 28 | 71 | 0.004765485 | 0.419 | -1.255 | 2.00E+00 |
| CHMP3 | 9 | 38 | 0.004855355 | 0.252 | -1.991 | 1.67E+00 |
| SPPL3 | 18 | 55 | 0.004855355 | 0.348 | -1.524 | 1.86E+00 |
| HLA-A | 17 | 1 | 0.004855355 | 18.061 | 4.175 | 1.26E+00 |
| IMMP2L | 698 | 591 | 0.004960151 | 1.255 | 0.328 | 3.11E+00 |
| PDAP1 | 6 | 31 | 0.00503364 | 0.206 | -2.282 | 1.57E+00 |
| RANBP3 | 116 | 67 | 0.005528212 | 1.839 | 0.879 | 2.26E+00 |
| ANKRD11 | 80 | 147 | 0.00588029 | 0.578 | -0.791 | 2.36E+00 |
| PHF12 | 52 | 107 | 0.006068487 | 0.516 | -0.954 | 2.20E+00 |
| SYNRG | 78 | 144 | 0.00634444 | 0.575 | -0.797 | 2.35E+00 |
| SRF | 11 | 41 | 0.006478703 | 0.285 | -1.811 | 1.72E+00 |
| LSM14A | 48 | 101 | 0.006702451 | 0.505 | -0.986 | 2.17E+00 |
| CDYL | 67 | 128 | 0.006790149 | 0.556 | -0.847 | 2.29E+00 |
| PDCD10 | 51 | 20 | 0.006790149 | 2.709 | 1.438 | 1.85E+00 |
| NFE2L1 | 118 | 198 | 0.006910871 | 0.633 | -0.660 | 2.50E+00 |
| CADM2 | 2377 | 2248 | 0.007029187 | 1.124 | 0.168 | 3.67E+00 |
| TAF1B | 5 | 28 | 0.007277261 | 0.190 | -2.398 | 1.52E+00 |
| SMURF1 | 127 | 210 | 0.007277261 | 0.642 | -0.638 | 2.53E+00 |
| COMMD7 | 29 | 71 | 0.007308639 | 0.434 | -1.205 | 2.00E+00 |
| CYLD | 71 | 34 | 0.007308639 | 2.219 | 1.150 | 2.02E+00 |
| SHMT2 | 31 | 75 | 0.007324311 | 0.439 | -1.187 | 2.03E+00 |

| | high.sense | low.sense | padjust.sense | MI.sense | log2MI.sense | log10 ins |
|---|---|---|---|---|---|---|
| SRRD | 1 | 18 | 0.007372275 | 0.059 | -4.083 | 1.28E+00 |
| XPOT | 42 | 91 | 0.007372275 | 0.490 | -1.028 | 2.12E+00 |
| RALGAPA2 | 307 | 232 | 0.007372275 | 1.406 | 0.492 | 2.73E+00 |
| EIF3K | 6 | 30 | 0.007511103 | 0.212 | -2.235 | 1.56E+00 |
| SPATA5 | 7 | 33 | 0.007533858 | 0.225 | -2.150 | 1.60E+00 |
| ASXL2 | 73 | 136 | 0.008204695 | 0.570 | -0.810 | 2.32E+00 |
| APAF1 | 48 | 100 | 0.008242648 | 0.510 | -0.972 | 2.17E+00 |
| GRIK2 | 846 | 739 | 0.008881607 | 1.216 | 0.283 | 3.20E+00 |
| FAM198B | 544 | 451 | 0.008881607 | 1.282 | 0.358 | 3.00E+00 |
| PRSS12 | 419 | 335 | 0.009053261 | 1.329 | 0.410 | 2.88E+00 |
| TBCK | 645 | 547 | 0.009367132 | 1.253 | 0.325 | 3.08E+00 |

**Supporting Information Table 3: HAP1.MR1 ATP13A1 CRISPR clones shown.** The change in sgRNA 2 ATP13A1 sufficient clones was necessary because clone 2-19 was bacterially contaminated in the set of experiments shown in Fig. 2. In experiments where both clones were used in parallel, comparable results were obtained. Like clone 3-1, clone 2-6 (but not clone 2-19) underwent morphological changes over time (see Supporting Information Fig. 7). The change in sgRNA 3 ATP13A1 sufficient clones was necessary because sequencing revealed gene editing of clone 3-15 in a highly conserved region of the protein that may impact functionality (see Supporting Information Fig. 6)

| | **sufficient** | | | **KO** | | |
|---|---|---|---|---|---|---|
| | sgRNA 1 | sgRNA 2 | sgRNA 3 | sgRNA 1 | sgRNA 2 | sgRNA 3 |
| Fig. 2 D-G | 1-17 | 2-6 | 3-2 | 1-7 | 2-20 | 3-1 |
| Fig. 5 A-C | 1-17 | 2-19 | 3-15 | 1-7 | 2-20 | 3-1 |
| SI Fig. 7 C | 1-17 | 2-19 | 3-15 | 1-7 | 2-20 | 3-1 |
| SI Fig. 8 | 1-17 | 2-19 | 3-15 | 1-7 | 2-20 | 3-1 |
| SI Fig. 9 C donor 32 | 1-17 | 2-19 | 3-2 | 1-7 | 2-20 | 3-1 |
| SI Fig. 10 | 1-17 | 2-19 | 3-15 | 1-7 | 2-20 | 3-1 |

**Supporting Information Table 4: Oligonucleotide sequences.** Adapter sequences for deep sequencing belong to Illumina, Inc. Oligonucleotide sequences © 2018 Illumina, Inc. All rights reserved. Derivative works created by Illumina customers are authorized for use with Illumina instruments and products only. All other uses are strictly prohibited. sgRNA sequences were designed based on the Sabatini library (1), TKO v3 (http://tko.ccbr.utoronto.ca), and GenomeCRISPR (http://genomecrispr.dkfz.de). XBP-1 primers have been published previously (2).

| | |
|---|---|
| **Linker for IRES Emerald removal** | |
| CK001_XhoI5_BamHI_NotI3 | TCGAGtgctagaaggcctacGGATCCGC |
| CK002_NotI5_BamHI_XhoI3 | GGCCGCGGATCCgtaggccttctagcaC |
| **Primers for the recovery of insertion sites** | |
| LTR sequencing primer | /doublebiotin/ggtctccaaatctcggtggaac |
| ssDNA linker for deep sequencing | /phospho/atcgtatgccgtcttctgcttgactcagtagttgtgcgat ggattgatg/dideoxycytidine/ |
| P5-containing adapter | AATGATACGGCGACCACCGAGATCTGATGGT TCTCTAGCTTGCC |
| P7-containing adapter | CAAGCAGAAGACGGCATACGA |
| sequencing primer | CTAGCTTGCCAAACCTAC AGGTGGGGTCTTTCA |
| **Primers for MR1-HA cloning** | |
| CK84_MR1BglII_fw | taaccgAGATCTccaccatgggggaactgatggcgttcc |
| CK85_MR1HANotI_rev2 | gctaaGCGGCCGCtcaAGCGTAATCTGGAACAT CGTATGGGTAtcgatctggtgttggaa |
| **Primers for XBP1 splicing assay** | |
| XBP1 splicing fw primer | ttacgggagaaaactcacggc |
| XBP1 splicing rev primer | gggtccaacttgtccagaatgc |
| **Plasmid sequencing** | |
| sffv1_promoterSeq_fw | TGCTTCTCGCTTCTGTTCG |
| CK082_MR1internal_fw | ctgcagtatgcatatgacgggcag |
| CK083_MR1out_fw | ctggagttggtgttctagtctg |
| **HAP1 CRISPR clone sequencing** | |
| CK24_ATP13A1_fw1 | acttcaggaccagtggctctgaa |
| CK25_ATP13A1_rev1 | ttcacaatgggcgtactctcgt |
| CK26_ATP13A1_fw2 | tcccagctttaagtgtggcacca |
| CK27_ATP13A1_rev2 | aaggacagcacctcaagcccgtcttc |
| CK28_ATP13A1_fw3 | gtggacttgtacaaggagtttgagccaa |
| CK29_ATP13A1_rev3 | acctgggcaatgaccagcttgaa |
| **THP-1 CRISPR clone sequencing** | |
| CK074_27illumina_rev2 | GTGACTGGAGTTCAGACGTGTGCTCTTCCGA TCTaaggacagcacctcaagcccgtcttc |
| CK80_79illumina_fw6 | ACACTCTTTCCCTACACGACGCTCTTCCGATC Tctgtaggttgagaagagctaagcc |
| Index D501 | AGGCTATA |
| Index D502 | GCCTCTAT |
| Index D503 | AGGATAGG |
| Index D504 | TCAGAGCC |
| Index D701 | ATTACTCG |
| Index D702 | TCCGGAGA |
| Index D703 | CGCTCATT |

| | |
|---|---|
| Index D704 | GAGATTCC |
| **Oligonucleotides for ATP13A1 CRISPR/Cas9 sgRNA cloning including BbsI adapters** | |
| guide1_sense | CACCGGGTTGTCGAGCGGCGACGG |
| guide1_antisense | AAATCCGTCGCCGCTCGACAACCC |
| guide2_sense | CACCGGTTGGCACCACCTTCACAA |
| guide2_antisense | AAATTTGTGAAGGTGGTGCCAACC |
| guide3_sense | CACCgGCAGGCTCTCCATGAAGGGC |
| guide3_antisense | AAATGCCCTTCATGGAGAGCCTGCc |
| **Target sequences for CRISPR/Cas9 sgRNA bulk validation** | |
| ATP13A1 sgRNA A | GGGTTGTCGAGCGGCGACGG |
| ATP13A1 sgRNA B | GCAGGCTCTCCATGAAGGGC |
| IRF2 sgRNA A | TAAACTCCAACACGATCCCG |
| IRF2 sgRNA B | CAGCGATGAAGAGAGTGCCG |
| NHLRC sgRNA A | TATCTGATGAGTCCCTGCCA |
| NHLRC sgRNA B | TCAGTACCCGAGTTTCCGGA |
| SPPL3 sgRNA A | GGCCAGTGAGAACCCAGATG |
| SPPL3 sgRNA B | GAGGCTCGGCAGGCGGACAA |
| VPS53 sgRNA A | GGAGGAACTGGAGTTCGTGG |
| VPS53 sgRNA B | TAAGAGGTCAGACGAACGTG |
| GANAB sgRNA A | GGAGCCAGCGAAGAAGGCCC |
| GANAB sgRNA B | TGTGGAGTTAACCATGGCTG |
| ZCCHC14 sgRNA A | GCATGGGAGCAGTGAGGCAA |
| ZCCHC14 sgRNA B | AAACCGACATACCTTCTCCA |
| TMEM131 sgRNA A | GGGACTCTGGCCCCAAGGAA |
| **Taqman probes used for qRT-PCR** | |
| HPRT | Hs02800695_m1 |
| ACTB | Hs01060665_g1 |
| GAPDH | Hs02758991_g1 |
| MR1 (routinely used) | Hs00155420_m1 |
| MR1 (alternative) | Hs01042278_m1 |
| ATP13A1 | Hs00220755_m1 |
| B2M | Hs99999907_m1 |

### Supporting Information References

1. Wang, T., Birsoy, K., Hughes, N. W., Krupczak, K. M., Post, Y., Wei, J. J., Lander, E. S., and Sabatini, D. M. (2015) Identification and characterization of essential genes in the human genome. *Science*. **350**, 1096–1101
2. Hollien, J., Lin, J. H., Li, H., Stevens, N., Walter, P., and Weissman, J. S. (2009) Regulated Ire1-dependent decay of messenger RNAs in mammalian cells. *Journal of Cell Biology*. **186**, 323–331
